# Molecular and functional basis of a novel Amazonian Dark Earth Esterase 1 (Ade1) with hysteresis behavior and quorum-quenching activity

**DOI:** 10.1101/2020.12.16.421545

**Authors:** Tania Churasacari Vinces, Anacleto Silva de Souza, Cecília F. Carvalho, Raphael D. Teixeira, Beatriz Aparecida Passos Bismara, Elisabete J. Vicente, José O. Pereira, Robson Francisco de Souza, Mauricio Yonamine, Sandro Roberto Marana, Chuck Shaker Farah, Cristiane R. Guzzo

**Author notes:** Corresponding author, and, To whom correspondence should be addressed: Prof. Cristiane Guzzo Carvalho, Department of Microbiology, Institute of Biomedical Sciences, University of São Paulo, Av. Prof. Lineu Prestes, 1374, Cidade Universitária, 05508-900, São Paulo-SP, Brazil.

## Abstract

Amazon Dark Earth (ADE) soil is rich in organic compounds and its fertility has been associated with a high diversity of microorganisms. Herein, we investigate the biochemical and functional features of a novel esterase, Ade1, obtained from a metagenomic library of Amazonian Dark Earth soils of the Amazonian Rainforest, in Brazil. The esterases cleave ester bonds to form a carboxylic and an alcohol group. Esterases and lipases are enzymes found in almost all living organisms, demonstrating their biological relevance. We reported that Ade1 belongs to an α/β-hydrolase superfamily. We suggest that Ade1 is a moonlighting enzyme with hysteresis behavior and quorum-quenching activity, which may play a key role in the metabolism of a Gram-negative proteobacteria. In addition, molecular dynamics simulations reveal that the hysteresis behavior is directly associated with structural properties of the cap domain. Our findings reveal details of the molecular basis, catalytic and structural mechanisms of a novel α/β-hydrolase, which may be applied to other esterases of biotechnological, food, and/or pharmaceutical interest.

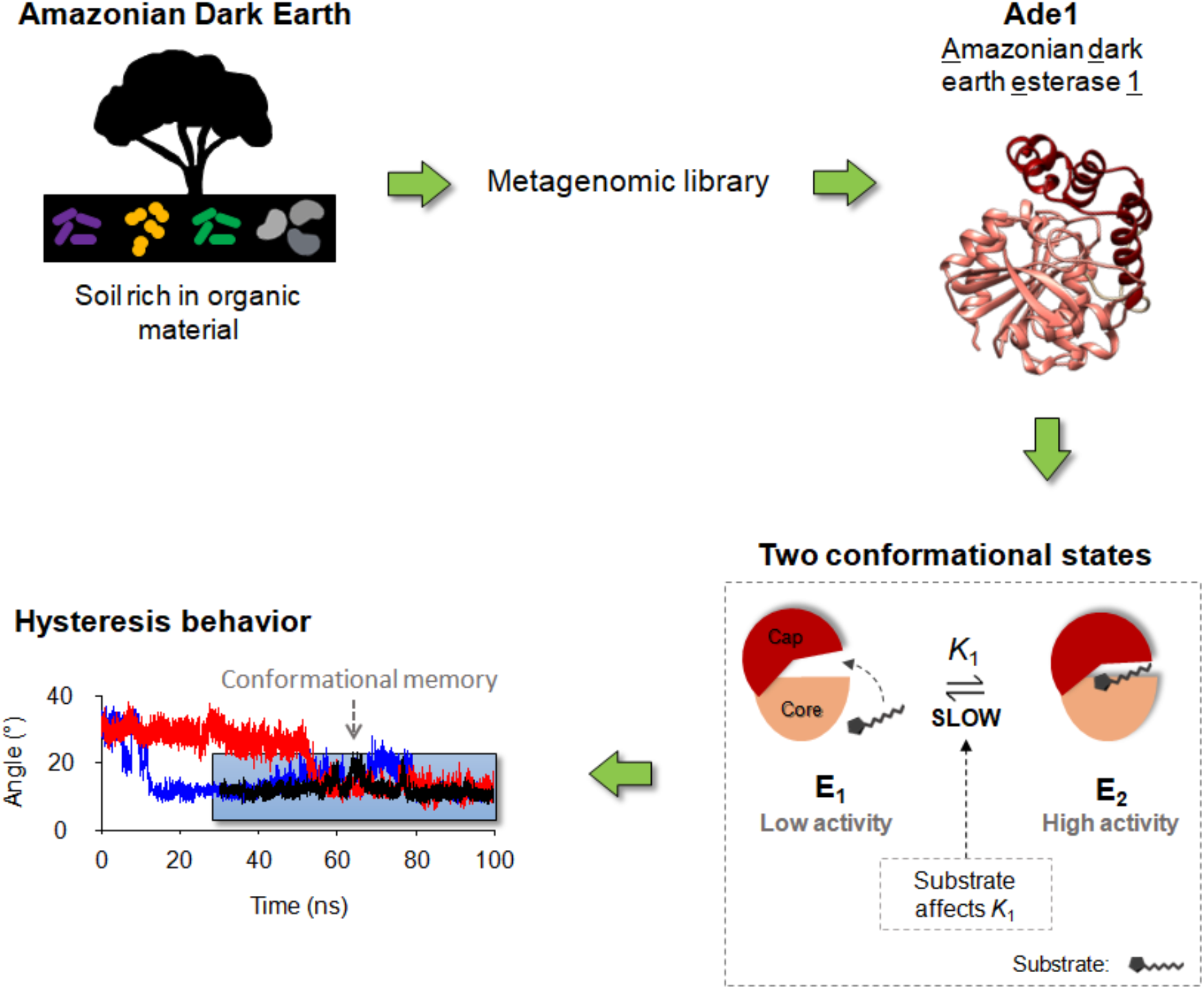

## Introduction

A particular type of soil, called Amazon Dark Earth (ADE), is characterized by their darker color, higher organic matter content, higher pH, and greater concentrations of phosphorus, calcium, magnesium, manganese, zinc, and other minor elements ^1^. These soils are of anthropogenic origin, derived from the activity of indigenous populations of the pre-Columbian era, and present a higher content of organic matter due to accumulation of vegetable and animal residues, in addition to a large amount of cinder and residues of bonfires (charcoal) ^2,3,42^. Its fertility is associated with the presence of high diversity of microorganisms ^5^. The composition and diversity of the microorganism communities in ADE soils have been explored and the most abundant bacterial phyla are *Proteobacteria* (24%), *Acidobacteria* (10%), *Verrucomicrobia* (8%), *Actinobacteria* (7%), *Planctomycetes* (4%), *Firmicutes* (3%), *Bacteroides* (3%), WS3 (2%), other less abundant clades (3%) as well as a considerable fraction of unclassified DNA samples (36%) ^32^. Within these microbial communities, distinct environmental factors select for a wide range of metabolic activities that are essential for the survival of microorganisms residing in these niches, each species possessing several enzymes that have the potential to be used in industry.

Arguably, for the classification of microorganisms in environmental samples, the metagenomic approach has gained prominence due to its potential for fast genetic analysis of all genomes contained in an environmental niche ^6^. In addition, it is a technique that provides genetic information for the discovery of biocatalysts or novel enzymes from uncultivated organisms, providing important insights about a wide diversity of a bacterial community ^7–10^. Relevant cell-cell communication is mediated by secretion of chemical compounds (autoinducer) to the environment, a process known as quorum-sensing (QS) ^11,12^. QS is activated when high cell density leads to high autoinducer concentration and cells respond with gene expression changes and coordinationated behavior ^13,14^. QS also regulates biofilm formation, antibiotic resistance, bioluminescence, and bacterial virulence ^12,15^. QS inhibition, known as quorum quenching (QQ), has revealed to play a fundamental physiological role for survival or microbial competition ^11,16,17^. One efficient way to impair cell-cell communication is by degradation or modification of an autoinducer by different classes of enzymes including lipolytic enzymes ^18,19^. QQ may have applications in the control of hospital and dental infections, plant diseases, and biofouling in the environment ^15^. In this context, N-acyl-homoserine lactones (AHLs) is one example of autoinducer well-characterized in Gram-negative bacteria ^11,12^.

Lipolytic enzymes are α/β-hydrolases characterized by the presence of a cap domain and a core, catalytic, domain ^20^. The core domain is described by an 8-stranded β-sheet surrounded by a variable number of α-helices ^20–22^, while the cap domain is composed by a variable number of α-helices. Enzymatic activity is due to the presence of a catalytic triad composed of a nucleophilic residue, a histidine and an acid residue ^22,23^. A nucleophile residue can be a serine, or cysteine, or an asparagine, while the acid residue can be an aspartate or glutamate ^22,23^. Lipolytic enzymes share the same physicochemical nature of catalytic triad, have high structure similarities and large potential for use in biotechnological processes, mostly due to their promiscuity, high regioselectivity, and stereoselectivity in different substrate reactions ^23^. To that end, lipolytic enzymes can be used in food modifications, detergent formulations, chemical synthesis, and drug metabolism and drug detoxifications ^11^.

Lipolytic enzymes are classified in two groups: esterase and lipases ^23^. Lipases are distinguished from esterases by the phenomenon of interfacial activation, in which the lipases hydrolyze long-chain esters (usually ≥ 10 carbons) ^23^, while esterases hydrolyze short-chain esters (usually < 10 carbons) ^23^. By having the same core structure and similar residues in the catalytic pocket,^22,23^ only experimental assays can help to identify their substrates and catalytic pathways. Esterases metabolize ester groups in alcohol and a carboxylic acid, and kinetic data tend to fit the Michaelis-Menten model ^24–29^. The Michaelis-Menten model assumes one or multiple independent active sites that bind one substrate and produce one product. Along the enzymatic turnover, the free enzyme conformation is restored for a new catalytic cycle^30^.

Interestingly, some monomeric enzymes show enzymatic activity with a profile of cooperativity and this characteristic is due to a slow conformational change in response to the substrate concentration, also known as hysteresis ^31–33^. There are two ways that cause enzymatic hysteresis behavior. One of them is the Mnemonic model, whose enzyme can keep a structural memory substrate-induced by a short time after releasing the product. In this model the enzyme visits only one conformational state in the absence of substrate. However, in the presence of substrate the enzyme visits two conformational states with different enzymatic activities. The other model that also causes hysteresis is the ligand-induced slow transitions (LIST). In this case, the enzyme visits two structural conformations in absence or presence of substrate. In contrast, the presence of substrate changes the equilibrium constant of the structural conformation states. In addition, the LIST behavior can be caused not only by the substrate but also by the solvent nature, pH, or applied voltage ^34–36^. Hysteresis is fundamental to understanding the system’s biology because its enzymatic profile can control activation or inhibition of a biological pathway using only one enzyme ^37^. One example of that is the modulation of the enzymatic activity based on the oscillation of a metabolic pathway benefiting the cell ^34–36^.

Herein, we characterized a novel esterase from a metagenomic library of Amazonian Dark Earths soil from the Amazonian Rainforest, in Brazil. We present its biochemical characterization as a novel esterase and, based on that, we named it as <u>A</u>mazonian <u>D</u>ark Earth <u>E</u>sterase <u>1</u> (Ade1). We solved its three-dimensional structure by crystallography, revealing an α/β-hydrolase fold and a catalytic triad composed by S94, D217, and H245. Kinetic data suggests that Ade1 has a hysteresis behavior contributing to the hypothesis that Ade1 is a metabolic enzyme. Molecular dynamics simulations of Ade1 confirmed a hysteresis behavior and associated it to structural properties of the cap domain. We also reported a quorum-quenching activity associated with AHL-producing cells. Ade1 is promiscuous and it is the first enzyme with a hysteresis behavior using p-nitrophenyl octanoate substrate and quorum-quenching activity.

## Results and Discussion

### Identification of a novel esterase enzyme from a metagenomic library

A metagenomic library was built from an Amazonian dark soil sample to search novel enzymes with esterase activities. To screen for diverse enzymatic activities, this library was built into fosmid vectors pCC1FOS with DNA fragments up to 40,000 bp and transformed into *E. coli* EPI300. After screening, 14 out of 80,000 colonies showed lipolytic activities, with formation of transparency halos due to hydrolysis of tributyrin. The best candidate was chosen to build sub-libraries to identify the gene responsible for the expression of lipolytic activity enzyme (Material and Method in Supporting Information). From the DNA sequencing, we identified an open read frame (ORF) of a gene whose length is 804 bp that translates to a protein of 268 residues (no signal peptide) and theoretical <u>m</u>olecular <u>w</u>eight (MW) of 29.3 kDa. We named it Ade1 (<u>A</u>mazonian <u>D</u>ark Earth <u>E</u>sterase <u>1</u>), whose nucleic acid and residue sequences are shown in **Table S1**.

In order to investigate the protein domains from our primary sequence and estimate its biological function, we submitted the translated ORF sequence to the Pfam web-based service at EBI. This analysis revealed that Ade1 belongs to the α/β-hydrolase superfamily ^38^ (*Abhydrolase_1* domain, Pfam PF00561). Comparing Ade1’s primary sequence to its best hit from the non-redundant NCBI protein sequence database, HOP18_27540 (GenBank: NOT58366.1), we observed 74% and 83% of amino acid sequence identity and similarity, respectively (**Figure S1**). Interestingly, HOP18_27540 was identified in contigs derived from DNA isolated in a metagenomic survey of drinking water samples from the Netherlands and assigned to the Deltaproteobacteria class ^39,40^ The high levels of sequence similarity to HOP18_27540 lead us to conclude that Ade1 is most likely encoded in the genome of a currently uncharacterized member of the Deltaproteobacteria class.

In order to classify the Ade1 family, we performed a phylogenetic tree analysis using 35 lipolytic enzyme families ^41^ (**Figure 1a**). We have found Ade1 belongs to family V and contains a conserved motif GXSXGG^42^ (**Figure 1b**). Members of family V are found in mesophilic bacteria (*Pseudomonas oleovorans, Haemophilus influenzae, Acetobacter pasteurianus*) as well as in cold-adapted (*Moraxella sp., Psy. immobilis*) or heat-adapted (*Sulfolobus acidocaldarius*) bacteria^43^. These members share primary sequence similarity between 20 to 25 % with non-lipolytic enzymes that include epoxide hydrolases, dehalogenases, and haloperoxidase, which also have an α/β-hydrolase fold ^44,45^.

**Figure 1.**
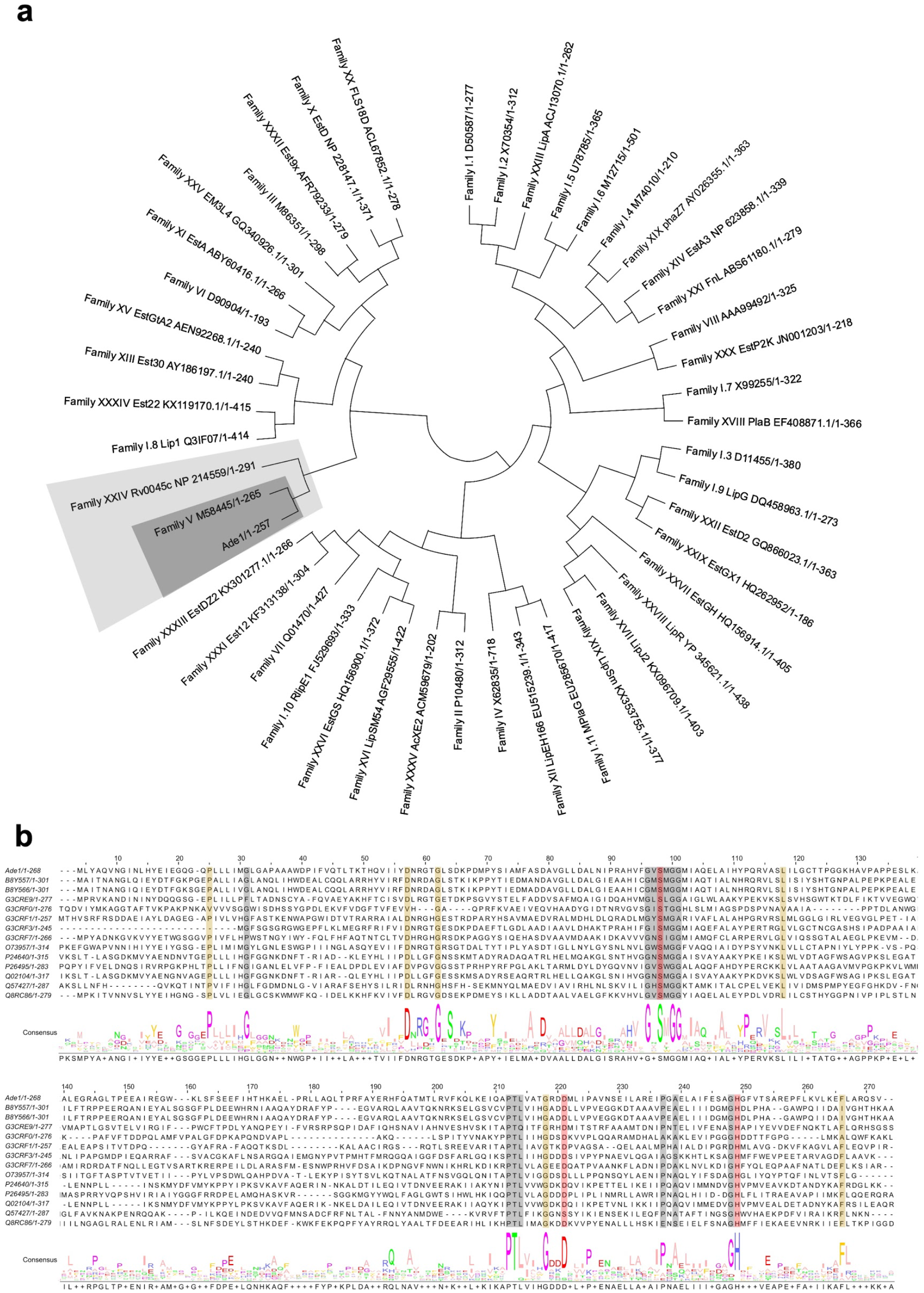
Classification and structural characterization of Ade1. **(a)** The polar tree layout representing the comparison analysis of the amino acid sequence of 35 families of lipolytic enzymes. It was constructed using the MEGA 10.1.8 program with the neighbor-joining algorithm. Each branch corresponds to one representative from each family with their respective UniProt ID. In the light gray color show the family groups that contained sequences more similar to Ade1; in dark gray, show the best candidate family would belong to Ade1, family V. **(b)** multiple sequence alignment of Ade1 and proteins belonging to family V. In red color, show the catalytic triad; in gray color, show the motif conserved only in the family V; and in yellow color, show other amino acids conserved. B8Y557: Lip 5 from *uncultured bacterium;* B8Y566: Lip 17 from *uncultured bacterium;* G3CRE9: AB hydrolase-1 domain-containing protein from the *uncultured organism*; G3CRF0: Peptidase_S9 domain-containing protein from the *uncultured organism*; G3CRF1: AB hydrolase-1 domain-containing protein from the *uncultured organism*; G3CRF3: AB hydrolase-1 domain-containing protein from the *uncultured organism*; G3CRF7: AB hydrolase-1 domain-containing protein from the *uncultured organism*; O73957: Sal from *Sulfolobus acidocaldarius*; P24640: Lip3 from *Moraxella sp*. (*strain TA144*); P26495: Poly(3-hydroxy alkanoates) depolymerase from *Pseudomonas oleovorans*; Q02104: Lip 1 from *Psychrobacter immobilis*; Q57427: Putative esterase/lipase HI_0193 from *Haemophilus influenzae*; the Q8RC86: Predicted hydrolase or acyltransferase from *Caldanaerobacter subterraneus subsp. tengcongensis*. In **Figure S17** is shown the dendrogram of the family V of lipolytic enzymes.

### Expression, purification, and structural characterization of Ade1

#### Cloning and expression

To determine the three-dimensional structure of Ade1, it was cloned in the expression vector pET-28a, transformed, and expressed into *E. coli* BL21(DE3)-RP cells. The soluble fractions were submitted in affinity chromatography to purify the Ade1 apo-state (Material and Method in Supporting Information and **Table S2**).

#### Structural characterization

In order to obtain the Ade1 holo-state, we added 4 mM tributyrin into purified Ade1 during the crystallization step. Protein crystals were submitted to X-ray diffraction and we obtained data up to 2.30 Å resolution. The phase was determined by molecular replacement using the crystal structure of an enol-lactone hydrolase (PcaD) from *Burkholderia xenovorans* (PDB ID 2XUA)^46^, which shares 29% and 44% of identity and similarity, respectively (**Figure S2**). Ade1 crystal diffraction data, processing, and refinement details are described in the Supporting information (Material and Method in Supporting Information and **Table S3)**. There are four Ade1 molecules in the asymmetric unit (ASU) presenting a root-mean-square deviation (RMSD) between C_α_ coordinates ranging from 0.1 to 0.2 Å. Ade1 structure has a typical α/β-hydrolase fold consisting of a core domain and a cap domain (**Figure 2**). Based on the chain B, the core domain is formed by one curved β-sheet containing 8 β-strands (from β1 to β8) surrounded by eight α-helices and each side of the β-sheet is surrounded by four α-helices (α1, α2, α12, α13 in one side and α3, α4, α10, α11 in the other side) (**Figures 2a** and **2b**). Cap domain (residues 121-196) has five α-helices (α5 to α9) (**Figures 2a** and **2b**). The main difference between the four Ade1 molecules in the ASU is in the loop between α5 and α6 located in the cap domain of Ade1. It is an unstructured region in chain A (residues 138-140) and in chain C (residues 138 - 143) (**Figure S3**). Comparing Ade1 with other members of α/β-hydrolases, we noted that the active site is located on the interface between the cap and the core domains, which contains the conserved catalytic triad, S94, D217, and H245 (**Figure 2a, c,** and **d**). The catalytic serine, S94, is buried inside the enzymatic pocket in a loop between α4 and β5 (known as a canonical nucleophile)^21^. Catalytic aspartate, D217, is localized in a loop connecting β5 and α11. Finally, the catalytic histidine residue is placed in a loop that connects β8 and α12. These localizations are fundamental for the enzymatic catalysis because they are involved with the closing of the cap and, afterward, reaction advance.

**Figure 2.**
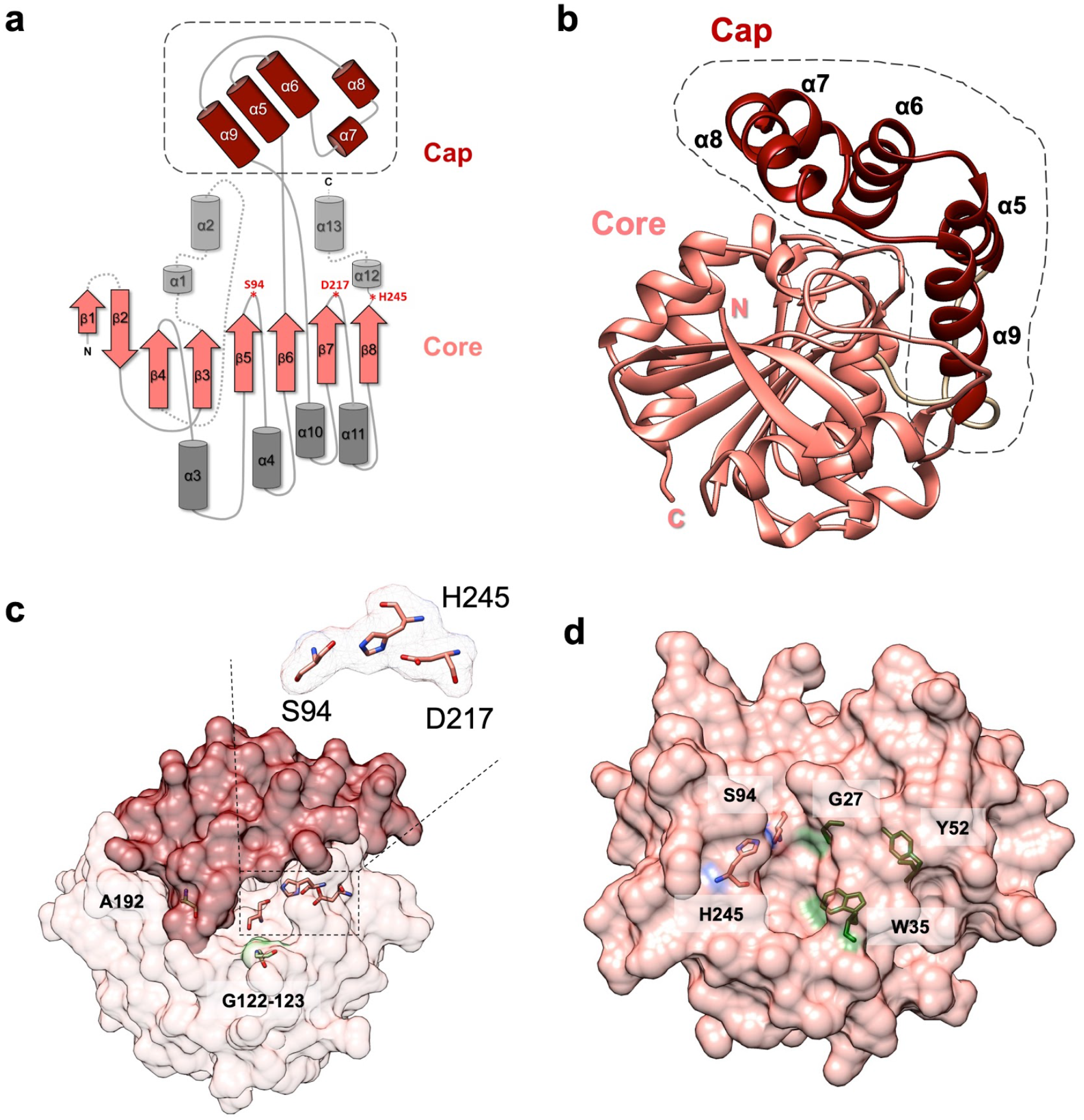
Ade1 structural characterization. Cap and core domains are colored of dark red and salmon colors. **(a)** Ade1 topology composed by eight β-sheet (arrows in salmon color) and thirteen α-helix (cylinder in gray and dark red colors), the cap (enclosed by dotted lines) is compound by α5-α8, the core or α/β hydrolase structural domain is compound by the total β-sheets connected by others α-helix (light and dark gray, behind and in front of β-sheets respectively), the catalytic triad S94, D217 and H245 show in red (asterisk). **(b)** 3D-structure of Ade1 represented in ribbon. **(c)** Ade1 represented by surface. It shows the catalytic triad S94, D217, and H245 (in salmon) and three high conserved residues A192, G122 and G123. **(d)** Core domain analysis, in which conserved amino acids are localized to the inside of the catalytic pocket. Besides, it shows the residues S94 and H245 (in salmon color) and G27, W35 and Y52 (in green color).

#### Conserved residues of Ade1

Multiple sequence alignment with Ade1 orthologous proteins identified conserved residues (numbering correspond to Ade1 protein and the bold residues are from the catalytic triad): Y_13_E_14_x2G_17_, G_19_xP_21_, G_27_, W_35_, Y_52_D_53_N_54_R_55_G_56_XG_58_, P_63_, Y_67_, A_73_xD_75_, A_88_x_3_G_92_x**S_94_**xG_96_G_97_x_2_A_100_Q_101_, L_114_xL_116_x_2_T_119_x_2_G_122_G_123_, A_192_, I_205_x_2_P_208_xL_210_x_3_G_214_x_2_**D_217_**, P_221_x_2_N_224_, A_229_x_2_I_232_, **H_245_**, and F_262_ (**Figure S4**). Mapping these conserved residues on the Ade1 structure, we observed that most of them are located in a core domain (**Figure S5**). However, the G122, G123, and A192 residues are located in the cap domain. G122 and G123 residues may be related to increasing the flexibility in the cap domain allowing an opening and closing of the cap domain. Interestingly, G17, G19, P21, G58, P63, P208, P221, and G123 are conserved and are exposed to the solvent. In addition, G27, W35, Y52, S94, and H245 residues are into the enzymatic pocket and are partially exposed to solvent (except S94, which is more exposed) (**Figure 2d** and **S5 c**). Interestingly, D217 is completely buried in the catalytic pocket, and helps to establish the H245 charge. S94 seems to be exposed to interact with the substrate and initiate the process of catalysis mediated by a nucleophilic attack by the serine. G27, W35, and Y52 also are conserved residues and may play a key role in the catalytic process.

#### Ade1 is in a monomeric state

Interestingly, analysis of protein-protein contacts using the <u>P</u>rotein <u>i</u>nterfaces, <u>s</u>urfaces, and <u>a</u>ssemblies (PISA) web-based service^47^ suggests that Ade1 is a monomer. Indeed, gel filtration coupled to the Multi-Angle Light Scattering detector (SEC-MALS) confirmed that Ade1 (with and without 6xHis-tag) is a monomer with an experimental MW of 30 kDa (theoretical MW of 29.3 kDa, **Figure S6**). Members of the α/β-hydrolase family also may oligomerize through β-strands and monomeric states are favored due to the presence of more than four α-helices in the cap domain ^48^. For example, PcaD has been described as a dimer in aqueous solution, in which the core domain (α1, β1, and β2) and its cap domain (α7) is involved in the protein-protein interaction ^46^. Therefore, structural features of the cap domain play a key role in the formation of oligomers ^49–51^. In agreement, the monomeric state of Ade1 is justified by the presence of five α-helices located in the cap domain.

#### Structural comparison between Ade1 and different α/β-hydrolase members

In order to verify the difference between Ade1 and α/β-hydrolase members, we compared the three-dimensional structures of them using DALI web-based service^52^. Ade1 presented detectable structural similarity with a wide range of α/β-hydrolase family members of different enzymatic functions. Among them are lactonases, peroxidases, lipases, epoxide hydrolases, peptidases, homoserine o-acetyltransferase, haloalkane dehalogenase, phospholipase, cutinase and different classes of esterases (such as arylesterase, acetylesterase, thioesterase, heroin esterase, cocaine esterase, acetylcholinesterase, sterol esterase, cholesterol esterase, carboxylesterases, and metallophore yersiniabactin (Ybt) protein)^53^ (**Table S4**).

We investigated the structural relationships between Ade1 and similar α/β-hydrolase structures using all-against-all comparisons in the DALI-server website (**Figure S7**, **Table S4**). **Figure S7a** shows a dendrogram representing a hierarchical clustering based on structure similarity scores between these structures. We identified 3 groups that do not share the same biochemical functions. Group 1 members have the same topology of Ade1 but present few structural differences in the cap domain (**Figure S7b**). Among group 2 members, we noted differences in the cap domain and insertions in the core domain relative to the structure of Ade1 (**Figure S7c**). Group 3 members are structurally more divergent from Ade1 and have significant differences in both the cap and core domain (**Figure S7d**). These data suggest that structural divergences among α/β-hydrolase superfamily members are not easily correlated to their different enzymatic activities. Three-dimensional structure in this family seems to be directly related to the structure and size of the substrate. As a consequence, for members of this superfamily, prediction of enzymatic activity and target substrates cannot be based solely on comparative analysis of the structures and biochemical assays are key for predicting the corresponding biological function of each α/β-hydrolase family member. In this context, we could not infer a specific enzymatic function for Ade1 from structure and chose, instead, to perform additional experiments to understand the biochemical and biological functions of the Ade1 ^52^.

### Functional characterization of Ade1

#### Biochemical studies

We evaluated the hydrolytic activity of Ade1 on tributyrin and triolein (**Figure 3a-b**). As a result, we did not detect a hydrolysis halo around the bacterial colony in the presence of triolein (**Figure 3b**). However, we observed a hydrolysis halo around the bacterial colony in the presence of tributyrin. Thus, we classified Ade1 as an esterase (**Figure 3a-b**). As expected, cells expressing the mutated Ade1 from catalytic serine to alanine, Ade1_S94A_, and cysteine, Ade1_S94C_, did not show hydrolytic activity in the presence of tributyrin (**Figure 3a**). We also evaluated the lipolytic activity of Ade1 through precipitation tests using Tween-20 (a linear substrate with aliphatic chain of 12 carbons) (**Figure 3c**) and Tween-80 (aliphatic chain of 17 carbons, data not shown). As a control, we used the Ade1_S94C_, which did not show lipolytic activity. Comparing the experimental conditions, we detected only the degradation of Tween-20 by observing the formation of insoluble white crystals due to the reaction of hydrolysis products with Ca^2+^. Taken together, Ade1 has esterase and tweenase activities.

**Figure 3.**
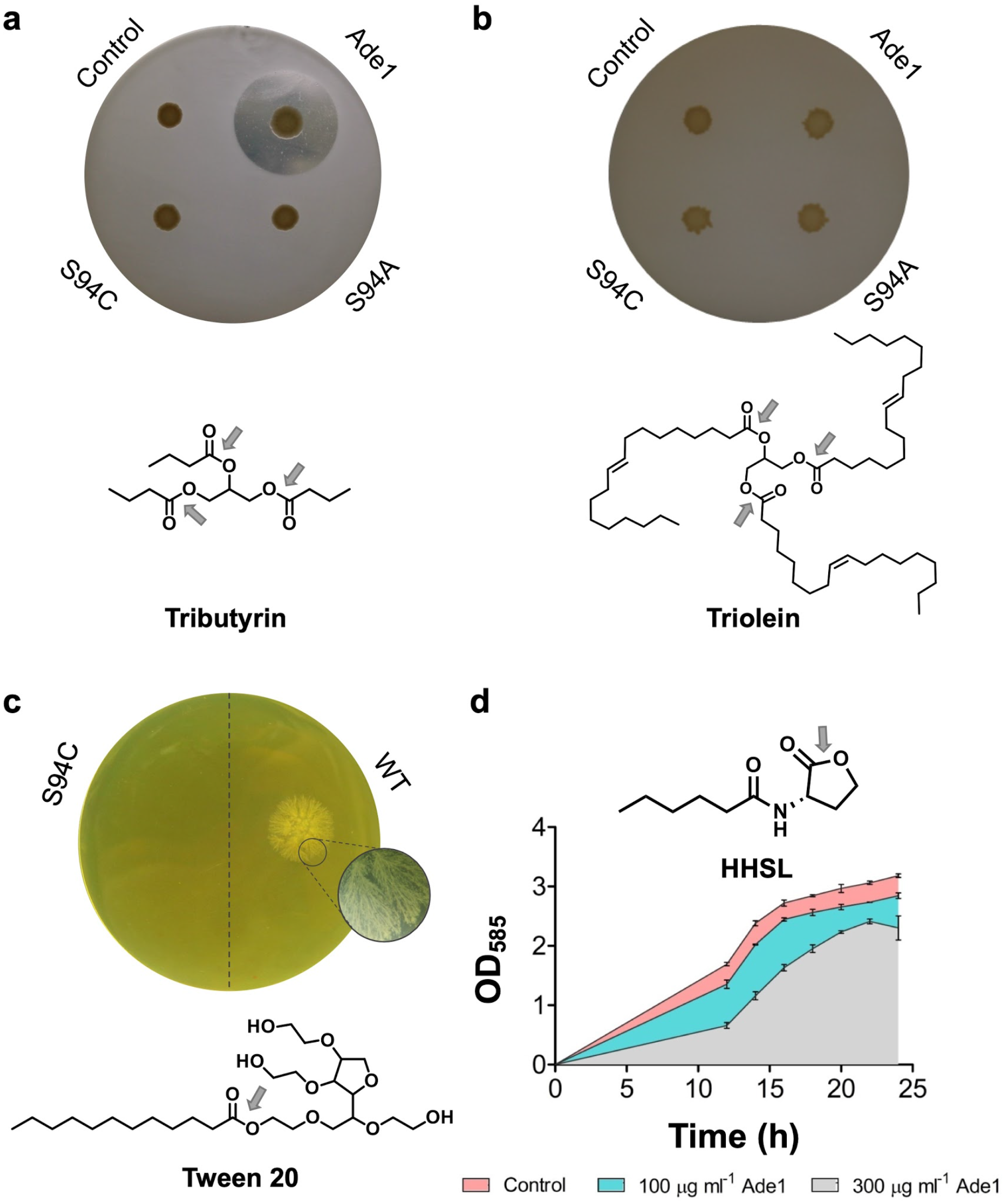
Esterase capacity of Ade1. **(a)** The hydrolysis of tributyrin due to *ade1* gene expression. Such hydrolysis generated a translucent halo around the colony suggesting that Ade1 is an esterase. **(b)** Ade1 does not hydrolysis triolein. **(c)** Tweenase activity tested with Ade1WT and Ade1_S94C_ after 24 hours. Insoluble crystals are observed around the protein inoculation because the reaction product can react with calcium. Inhibition of violacein production due to presence of Ade1. **d**) Violacein production was measured at O. D. 585 nm, which is an indirect relation of the HHSL concentration. *Chromobacterium violaceum* growth in absence (salmon color) and presence of 100 μg.ml^-1^ and 300 μg.ml^-1^ of Ade1 (ciane and gray colors, respectively), in which O. D. 585 nm. Each assay was performed in triplicates. Grey arrows indicate the hydrolysis sites.

Ade1 has significant structural similarity to cocaine esterase (**Figure S7d** and **table S4**), an enzyme that hydrolyzes pharmacologically active cocaine to a nonpsychoactive metabolite^54^. We therefore tested Ade1’s capacity to hydrolyze the cocaine molecule, a substrate that contains a 8-methyl-8-azabicyclo[3.2.1]octane and a benzyl group. We did not observe significant hydrolysis of cocaine to ecgonine methyl ester and benzoic acid or benzoylecgonine (data not shown). This result suggests that Ade1 is not able to hydrolyze esters connected to steric bulky groups.

#### Quorum-quenching activity

Dimeric enol-lactone hydrolase (ELH) PcaD (expressed by *Burkholderia xenovorans)* cleaves the lactone ring of 3-oxoadipate-enol-lactone (also known as β-ketoadipate-enol-lactone) to β-ketoadipate ^46^. This molecule is converted to succinate and acetyl-CoA, which are intermediates of the Krebs cycle via β-ketoadipyl-CoA ^55^. PcaD is an esterase involved in the bacterium metabolism by the degradation of aromatic compounds (protocatechuate or catechol) to use the products on the KREBS cycle to obtain energy. It can be used to degrade homoserine lactones by blocking quorum-sensing as a biotechnological tool ^46^.

Since Ade1 shares 44% primary sequence similarities and high structural similarity with PcaD (RMSD of 2.3 Å, **Table S4**), we performed an assay for detecting whether Ade1 has quorum-quenching activity. To that end, we monitored the growth of the *Chromobacterium violaceum* that produces a characteristic purple pigment, known as violacein, in a quorum sensing (QS) dependent way. Synthesis of violacein is activated in the presence of *N*-<u>h</u>exanoyl-L-<u>h</u>omo<u>s</u>erine <u>l</u>actone (HHSL), which activates the QS and biosynthesis of violacein ^56^. Thus, measurement of violacein pigment is an indirect method for detecting the HHSL. Ade1 concentrations from 100 to 300 μg.ml^-1^ in the medium culture decrease significantly violacein production by *Chromobacterium violaceum* (the area under the curve is decreased with an increase of the Ade1 concentration) (**Figure 3d**). Therefore, our results suggest that Ade1 has quorum-quenching activity by cleaving HHSL since Ade1 in the concentrations used do not affect the *Chromobacterium violaceum* growth curve (data not shown).

Quorum-quenching by degradation of homoserine lactones is observed in different classes of enzymes such as lactonases, metallo-β-lactamases, decarboxylase, acylase, oxidoreductase and deaminase (see more information in the review ^57^). These enzymes cleave the homoserine lactone at different sites of the homoserine lactone and may have different mechanisms of catalysis. We analysed some enzymes with quorum-quenching activity and most of them are described as esterases (**Table S5**). Few of them have been reported to have their enzymatic activity affected by the presence of bivalent or monovalent metals. However, metallo-β-lactamases require two Zn^2+^ or Co^+2^ ions for enzymatic activity ^58,59^ (**Table S5**).

### Kinetic activity of Ade1

#### Enzymatic kinetic studies

In order to study the esterase kinetic activity of Ade1, we performed enzymatic assays using a different substrate containing *p*-nitrophenyl ester (*p*-NP) with different aliphatic group sizes (from 4 to 16 carbons). Reaction product, *p*-nitrophenyl, was measured by absorbance at 347 nm. Initially, we evaluated the Ade1 activity in the presence of *p*-nitrophenyl butyrate, *p*-nitrophenyl octanoate, *p*-nitrophenyl laurate, and *p*-nitrophenyl palmitate (**Figure 4**). Ade1 hydrolyzes only *p*-nitrophenyl butyrate and *p*-nitrophenyl octanoate and we do not detect significant enzymatic activity using *p*-nitrophenyl laurate and *p*-nitrophenyl palmitate. Thus, substrates with aliphatic carbon chains containing more than 11 carbons did not present detectable reaction products. Interestingly, Ade1 preferentially catalyzes the hydrolysis of *p*-nitrophenyl octanoate (**Figure 4**). In addition, we tested p-nitrophenyl octanoate as substrates for Ade1_S94A_ and Ade1_S94C_ but both mutants did not present detectable enzymatic activities (**Figure S8**).

**Figure 4.**
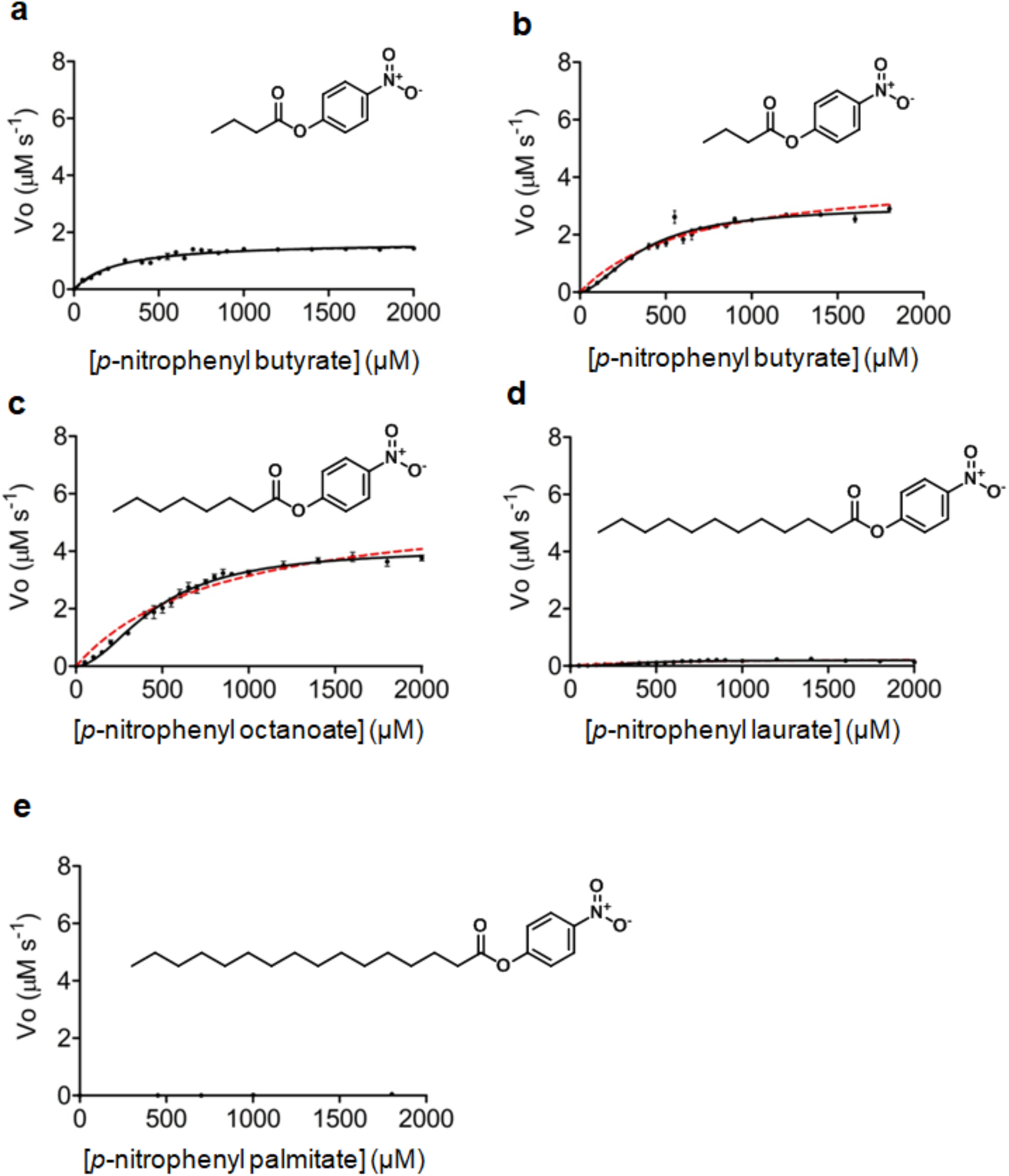
Enzymatic characterization. Kinetic activity of Ade1 in presence of different substrates. p-nitrophenyl butyrate in **(a)** presence and **(b)** absence of Triton X-100. **(c)** p-nitrophenyl octanoate in presence of Triton-X100. Hydrolysis of **(d)** p-nitrophenyl laureate, **(e)** p-nitrophenyl decanoate and **(f)** p-nitrophenyl palmitate, all without Triton X-100. In panels **(b)** and **(c)** are shown two curves corresponding to Michalis-Menten (dotted lines, red color) and Hill-Langmuir (continuous lines, black color) models. In all kinetics assays (with exception *p*-nitrophenyl butyrate), the hypothesis test showed that Hill-Langmuir model best represents the experimental data (*p*-value less than 0.001).

In order to determine the optimal enzymatic activity conditions of Ade1, we performed different kinetics assays using *p*-nitrophenyl octanoate as a substrate. We measured the dependence of esterase activity on the protein concentration ranging from 20 to 80 nM (**Figure S9**). As expected, we did not detect significant differences in the specific activity of Ade1. Next, we investigated the effect of pH on the reaction velocity. Ranging pH from 4.0 to 9.0, we noted that the maximum enzymatic reaction velocity was reached in the range of 7.0 to 9.0 (**Figure S10a**). We also evaluated if bivalent metals Ca^2+^, Co^2+^, Mg^2+^, Ni^2+^, and Zn^2+^ may affect the enzymatic activity. Curiously, increasing the Co^2+^ concentration from 0 to 3 and 6 mM, we observed that reaction velocity increased from 0.75 to 2.27 and 3.7 μM.s^-1^, respectively (**Figure S10b**). Therefore, Co^2+^ improves Ade1 catalytic activity. We observed that the presence of Zn^2+^ abolished Ade1’s activity and caused protein aggregation (data not shown). It is impressive how the enzymatic activity of Ade 1 is unaffected by high concentrations of bivalent cations (concentrations ranging from 1 to 6 mM) and the enzyme significantly increases its enzymatic activity by 6 mM cobalt.

Interestingly, we observed a hyperbolic curve using *p*-nitrophenyl butyrate (**Figure 4a**) and a sigmoidal profile curve using *p*-nitrophenyl octanoate as substrates (**Figure 4c**). We also performed the same assay using *p*-nitrophenyl butyrate as a substrate in the absence of Triton-X100 and a sigmoidal profile became more evident (**Figure 4b** compared to **figure 4a**). Intriguingly, the presence of Co^2+^ in the reaction buffer using as a substrate *p*-nitrophenyl butyrate or *p*-nitrophenyl octanoate causes a sigmoidal curve more prominent (**Figure 5 a-d**). This profile is not expected because Ade1 is a monomer. Cooperative behavior would be observed if Co^2+^ induced Ade1 oligomerization. Therefore, we performed SEC-MALS assays in the presence of Co^2+^ and we did not observe any oligomeric state for Ade1 (**Figure S6**). Nevertheless, this result suggests that Co^2+^ may cause small conformational changes, as suggested by the presence of a tiny shift in the SEC-MALS chromatography curve (**Figure S6**). In addition, the hydrolysis of *p*-nitrophenyl octanoate demonstrated a sigmoidal profile in absence and presence of Co^2+^ (**Figure 5c-d**). Increasing Co^2+^ concentration from 0 (presence of EDTA) to 0.2 and 6 mM, affects the velocity of hydrolysis of *p*-nitrophenyl octanoate. Furthermore, the sigmoidal curve, as described before, is more evident when we compare 0.2 and 6 mM Co^2+^ to the condition without this metal (**Figure 5e**).

**Figure 5.**
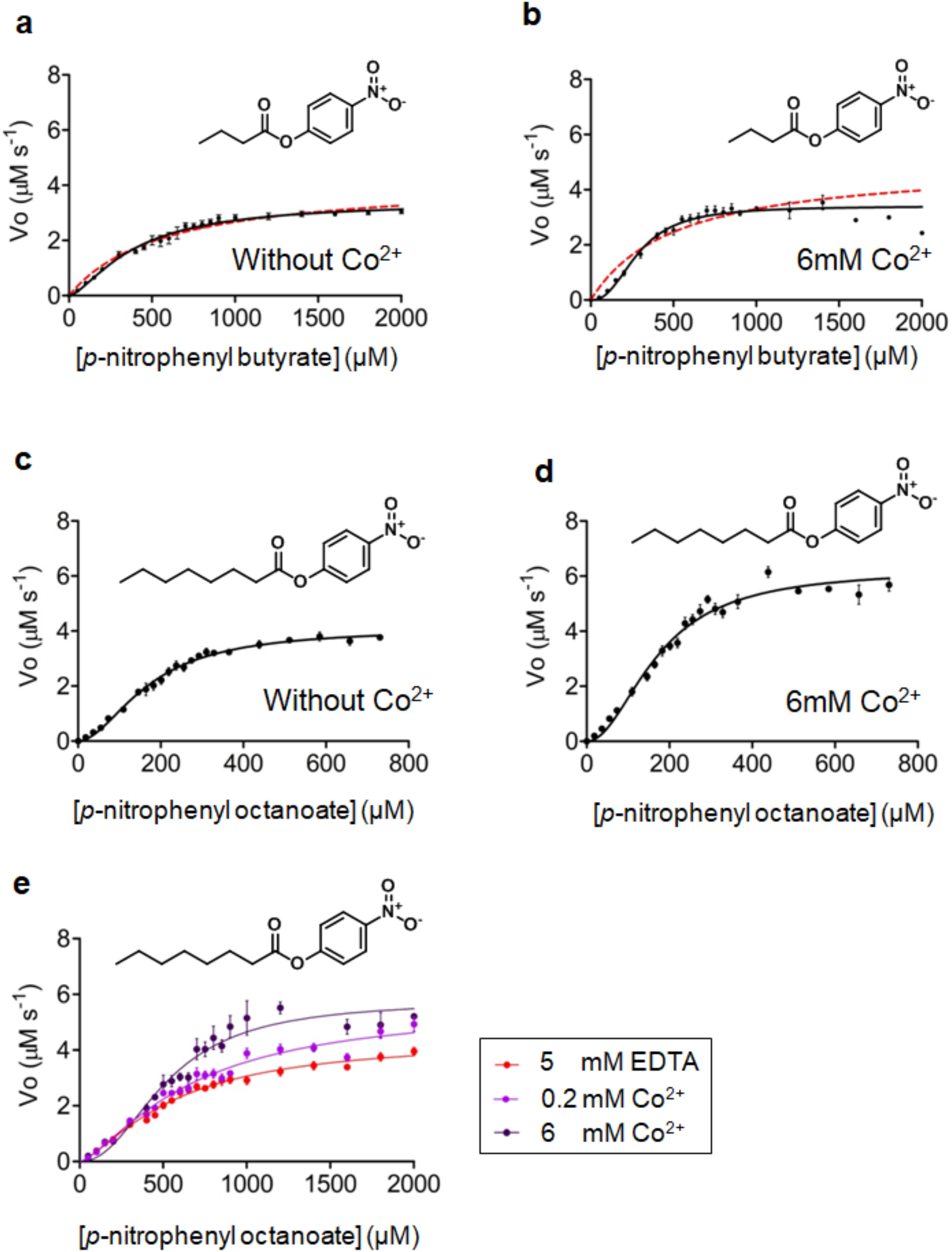
Hydrolysis of substrates p-nitrophenyl butyrate and *p*-nitrophenyl octanoate. Kinetic activity of Ade1 in presence of different substrates. *p*-nitrophenyl butyrate in **(a)** absence and **(b)** presence of 6 mM Co^2+^. *p*-nitrophenyl octanoate in **(c)** presence and **(d)** absence of 6 mM Co^2+^. In all kinetics assays (with exception *p*-nitrophenyl butyrate), the hypothesis test showed that Hill-Langmuir model best represents the experimental data (*p*-value less than 0.001). **(e)** Initial reaction velocity of *p*-nitrophenyl octanoate at different Co^2+^ concentration (0 mM, red color; 0.2 mM, purple; 6 mM, black). In panels **(a)** and **(b)** are shown two curves corresponding to Michalis-Menten (dotted lines, red color) and Hill-Langmuir (continuous lines, black color) models.

We initially hypothesized that the sigmoidal profile of reaction velocity could be caused by the presence of Triton X-100 in the reaction buffer, in which the substrate would be in equilibrium between a free state in solution and inserted into the Triton X-100 micelles. Thus, the enzyme would bind only free aggregates of substrate in solution. Based on that, we proposed a kinetic scheme shown in **Figure S11a**. We called this hypothesis the Substrate Effect model. Initially we detected the presence of free *p*-nitrophenyl octanoate in solution (**Figure S12**) in order to observe if when plotting initial velocities versus free substrate state we would observe a hiperbolic profile. Nevertheless, concentration of free substrate followed a direct relationship with the total concentration of *p*-nitrophenyl octanoate present in the enzymatic reaction (**Figure S12d**) contrasting with the substrate effect model which proposed a non-linear profile. Indeed it shows that ~36.5% of the total substrate is free in solution (**Figure S12**). Therefore, these observations invalidate the Substrate Effect model. The presence of Triton X-100 is not causing the sigmoidal effect. This conclusion is also supported by the observation that the hydrolysis of *p*-nitrophenyl butyrate in the absence of Triton X-100 also has a slight sigmoidal curve profile in plots of the initial velocities versus substrate (**Figure 4 a-b**). Moreover, when we performed the same assay without Triton X-100, but in the presence of Co^2+^, the sigmoidal curve is more evident (**Figure 5 a-b**).

Another explanation for a sigmoidal profile for Ade1 enzymatic activity would be the presence of hysteresis for the monomeric forms of the enzyme (**Figure S11 b**). In this model, the enzyme is in equilibrium (K1) between two conformational states, E1 and E2, in which the K1 is affected by substrate concentration. These states have different activities due to their different conformational states. Kinetics data of hysteretic enzymes are adequately described by the Hill-Langmuir model ^60–62^.

We performed an extra sum-of-squares F test in the GraphPad PRISM program to compare the fits using Michaelis-Menten and Hysteresis models. Data fit significantly better with Hysteresis models (Hill-Langmuir model), suggesting that Ade1 has hysteresis behavior (**Figure 4 b-c, Figure 5 a-d**, *p*-value < 0.001). Based on that, we fitted our data using a hysteresis model and determined the Ade1 kinetics parameters (**Table 1**). One exception is when the substrate is *p*-nitrophenyl butyrate in the presence of Triton X-100 in the reaction buffer, conditions where the kinetics data fits better with Michaelis-Menten model (**Figure 4a** and **Table 1**). In the case of *p*-nitrophenyl octanoate, the kinetics parameters were obtained in the presence and absence of Co^2+^ considering free *p*-nitrophenyl octanoate concentration instead of total concentration (**Figure 5 c-d**).

**Table 1.** Enzymatic kinetic data of Ade1. Kinetic parameters calculated from **Figures 4 - 5** using Hill-Langmuir or Michalis Menten model. The enzymatic assays were performed using 36 nM of Ade1. ***K*_half_** is similar to Michaelis-Menten constant (*K*_M_).

| Substrate | $V_{max}$ ($\mu M \cdot s^{-1}$) | $K_{half}$ ($\mu M$) | $k_{cat}$ ($s^{-1}$)<sup>#</sup> | $k_{cat}/K_{half}$ ($\mu M^{-1} s^{-1}$) |
| --- | --- | --- | --- | --- |
| *p*-nitrophenyl butyrate<sup>&</sup> | $3.4 \pm 0.1$ | $392 \pm 29$ | $94 \pm 3$ | $0.24 \pm 0.03$ |
| *p*-nitrophenyl butyrate<sup>*&</sup> | $3.4 \pm 0.1$ | $288 \pm 12$ | $94 \pm 3$ | $0.33 \pm 0.02$ |
| *p*-nitrophenyl butyrate<sup>!</sup> | $1.7 \pm 0.1$ | $254 \pm 25$ | $47 \pm 1$ | $0.18 \pm 0.02$ |
| *p*-nitrophenyl butyrate<sup>*</sup> | $3.0 \pm 0.2$ | $377 \pm 30$ | $83 \pm 6$ | $0.22 \pm 0.03$ |
| *p*-nitrophenyl octanoate | $4.1 \pm 0.1$ | $171 \pm 7$ | $114 \pm 3$ | $0.67 \pm 0.04$ |
| *p*-nitrophenyl octanoate<sup>*</sup> | $6.2 \pm 0.3$ | $171 \pm 7$ | $172 \pm 8$ | $1.01 \pm 0.09$ |

\* added 6mM $Co^{2+}$
& Without triton X-100 in the reaction buffer
<sup>#</sup> $k_{cat} = V_{max}/[E]_{total}$
! Calculated based on Michaelis-Menten equation

Interestingly, the presence of Co^2+^ did not affect the *V*_max_ of *p*-nitrophenyl butyrate (3.4 ± 0.1 μM.s^-1^) but *K*_half_ decreased from 392 ± 29 to 288 ± 12 μM. These results suggest that Co^2+^ increases Ade1 substrate binding affinity (**Figure 5, Table 1**). On the other hand, when the reaction buffer has Triton X-100, the presence of Co^2+^ increases significantly the *V*_max_ of *p*-nitrophenyl butyrate hydrolysis from 1.68 ± 0.04 to 3.0 ± 0.2 μM.s^-1^, while the *K*_half_ increases from 254 ± 25 to 377 ± 30 μM. These results suggest that Co^2+^ decreases Ade1 substrate binding affinity in the presence of Triton X-100 (**Figure 5, Table 1**).

Considering the *p*-nitrophenyl octanoate hydrolysis in the presence and absence of Co^2+^, we observed that the *V*_max_ increased from 4.1 ± 0.1 to 6.2 ± 0.3 μM.s^-1^, respectively (**Figure 5, Table 1**). Nevertheless, *K*_half_ presented the same value at 171 ± 7 μM, demonstrating that the binding affinity is not affected. Therefore, in all experimental conditions tested, Co^2+^ increases the Ade1 catalytic efficiency (*k*_cat_/ *K*_half_, **Table 1**) probably by causing a conformational change to a higher activity state.

Ade1 has a high catalytic efficiency for *p*-nitrophenyl octanoate in the presence of Co^2+^ (1.0 μM^-1^s^-1^) when compared with the other *p*-NP substrates tested (**Table 1**). Nevertheless, its catalytic efficiency is low when compared to carboxylesterase PestE from a hyperthermophilic archaeon *Pyrobaculum calidifontis* (31.4 μM^-1^s^-1^ for *p*-NP caprylate (C8) ^63^). Nevertheless, Ade1 has high catalytic efficiency when compared with other bacterial and fungal esterases ^64^, such as RmEstA and RmEstA from fungal *Rhizomucor miehei* (0.5 10^-3^ and 4 10^-6^ μM^-1^s^-1^ of catalytic efficiency for *p*-NP caprylate (C8), respectively ^65,66^), Est5 from a metagenomic library (0.16 μM^-1^s^-1^ of catalytic efficiency for *p*-nitrophenyl octanoate ^67^) and EstOF4 from *Bacillus pseudofirmus* OF4 (0.02 μM^-1^s^-1^ of catalytic efficiency for *p*-NP caprylate ^68^).

#### Experimental data show that Ade1 has a hysteresis behavior

Hysteretic enzymes respond to substrate concentration change, which plays a key role in bacterial metabolic regulation ^69^. It has been reported that enzymes with hysteretic behavior can be dependent on both the structural aspects and the chemical nature of the substrate ^70^. It can be manifested as transient burst or lag. In burst, it exhibits a high initial reaction velocity and then decreases and stabilizes the velocity rate; the lag exhibits low initial reaction velocity, which after a time span, increases and stabilizes ^71^. **Table S7** shows the insights about different enzymes with hysteresis behavior, transient types (lag or burst). Thus, we noted that only 4 enzymes were attributed as a transient burst and 9 enzymes were characterized as transient lag. In general, hysteresis behaviors of these enzymes are activated by temperature, metals, pH, or substrate structure, and are directly associated with metabolic regulation. Interestingly, we observed that Ade1 presents a transient burst profile along with the hydrolysis of the substrates *p*-nitrophenyl butyrate and *p*-nitrophenyl octanoate (**Figure S13**). Therefore, we proposed that Ade1 exhibits two conformational states, E1 and E2, in which the E2 state is more active than E1. Substrate concentration affects the E1 and E2 equilibrium constant during the course of enzymatic reaction. Therefore, during reaction catalysis E2 is in higher concentration when the substrate is in excess and E2 concentration decreases as the substrate concentration declines. This mechanism of autoregulation by the substrate concentration may be an efficient way to modulate a metabolic pathway and can be important for the cell growth fitness.

### Computational simulation studies

#### Structural analysis of the apo- and holo-states of Ade1

In order to understand the structural changes during the Ade1 enzymatic process, we performed a molecular dynamics simulation of the Ade1 apo- and holo-states (using tributyrin as substrate) (**Figure S14**). **Figure S14a** shows the root-mean-square fluctuation (RMSF) of both states of Ade1. Thus, we identified the residues 123-206 as a critical region, which involves predominantly the cap domain (residues 121-196). Interestingly, we have seen that this region is more flexible in Ade1 apo-state than in the holo-state (**Figure S14a**). Substrate seems to induce a structural stability in the cap domain favoring the closed conformation.

We noted that this high flexibility is associated with an opening and closing of the cap domain (enzymatic cavity) as shown by calculating the angle formed by residues V127-L22-V194 (**Figure S14b**). Ade1 apo-state presents the cap domain opened (~30°) from 0 to ~50 ns (corresponding to 50% of the time along MD simulation) (**Figure S14c**), which is directly associated with low conformational changes (**Figure S14d**). Afterward, Ade1 apo-state maintains its cap domain closed (~15°) from ~50 to 100 ns (**Figure S14c**), which is related to high conformational changes of the enzyme (**Figure S14d**). Conversely, Ade1 holo-state presents the cap domain in the opened conformation during ~10 ns and then after that, it is closed along the remaining ~90 ns of the MD (**Figure S14e**). Such behaviors are correlated with high conformational changes of Ade1 (**Figure S14e**). These analyzes prompt us to define two conformational states, which are associated with the opening (E1 state) and closing (E2 state) of the cap domain. Interestingly, it has been described that MenH involved in the vitamin K biosynthetic pathway, an α/β-hydrolase, also has the same behavior of closing and opening of the cap domain ^72^. Conformational transition from open (low activity conformation) to closed (high activity conformation) involves movement of the cap domain ^72^. In the cap closed conformation, the side chain of catalytic histidine residue changes from a disfavored position to form a functional catalytic triad ^72^. The functional triad is activated when the histidine is close to the nucleophile serine and activates it to attack the substrate. We suggest that Ade1 is in an equilibrium between E1 and E2 states in both conditions, absence and presence of substrate. Nevertheless, the presence of substrate changes this equilibrium constant, *K*_1_, favoring the E2 state, which is consistent to a hysteresis behavior (**Figure S13** and **S14**).

#### Catalysis mechanism is associated with hysteresis behavior

We investigated hydrogen bonding interactions between the side chains of the S94 and H245 residues with nearby residues inside the enzymatic cavity along molecular dynamics simulation trajectories (**Table S8**). Interestingly, the occupancies of hydrogen bond between the pairs H245/D217 and S94/H245 presented values of 69.3 and 13.5 %, respectively. These results suggest that the interaction between H245 and D217 is key for stabilizing the interaction between S94 and H245. Considering these results, we combined our simulation results with a proposed catalysis mechanism for the cocaine esterase ^73^ to predict the Ade1 catalytic mechanism (**Figure S15**). Initially, the cap domain is open and receives the substrate. The reaction mechanism has two stages, (I) acylation and (II) deacylation (**Figure S15**). In the acylation stage, the D217 plays an important role increasing the basicity of the H245. Thus, a proton is transferred from S94 to H245 to form an ionic pair (S94^-^/H245^+^), which plays a critical role in the enzymatic reaction. Ionic pair formation may be favored also due to a salt bridge interaction between the residues D217 and H245. After the ionic pair formation, S94^-^/H245^+^, the charged serine makes a nucleophilic attack on the carbonyl carbon of the substrate to produce a transient tetrahedral intermediate (TTI). Oxygen 2 electron pair forms a π bond and the H245 residue donates a proton HE2 to oxygen 1 of TTI, forming the reaction alcohol product, R_1_-OH. In the deacylation stage, a base-catalyzed reaction is carried out, in which H245 acts as a Lewis base activating a water molecule to make a nucleophilic attack. The water molecule attacks the carbonyl carbon of the intermediate state of S94 bound covalently to substrate to form a novel TTI. Finally, the oxygen 2 electron pair forms a π bond, and the initial state of the catalytic serine is restored by transfer of H245 HE2 to S94 and the carboxylic acid, R2–COOH, is released.

Next, according to our experimental data, we considered in our mechanistic proposal, an enzymatic catalysis combinated with the hysteresis behavior of Ade1 (**Figure 6a-b**). To verify this biochemical feature in a structural context, we removed the tributyrin of the Ade1 holo-state after 30 ns of simulation to observe if the protein does not restore to the E1 state (apo conformation) and keeps the conformational state memory E2 (hysteresis simulation, **Figure 6b**). Comparing the backbone RMSD of the Ade1 apo- and holo-states with the hysteresis simulation, we observed a structural behavior more similar to Ade1 holo-state than Ade1 apo-state (**Figure 6c**) and it is more similar to closed than to open conformation (**Figure 6d**). Interestingly, the rootmean-square fluctuation (RMSF) also shows that (**Figures 6e-f**) hysteresis simulation is more similar to Ade1 holo-than apo-state profile. It suggests that Ade1 keeps a structural memory of the Ade1 closed conformation after substrate binding and product release consistent to hysteresis behavior.

**Figure 6.**
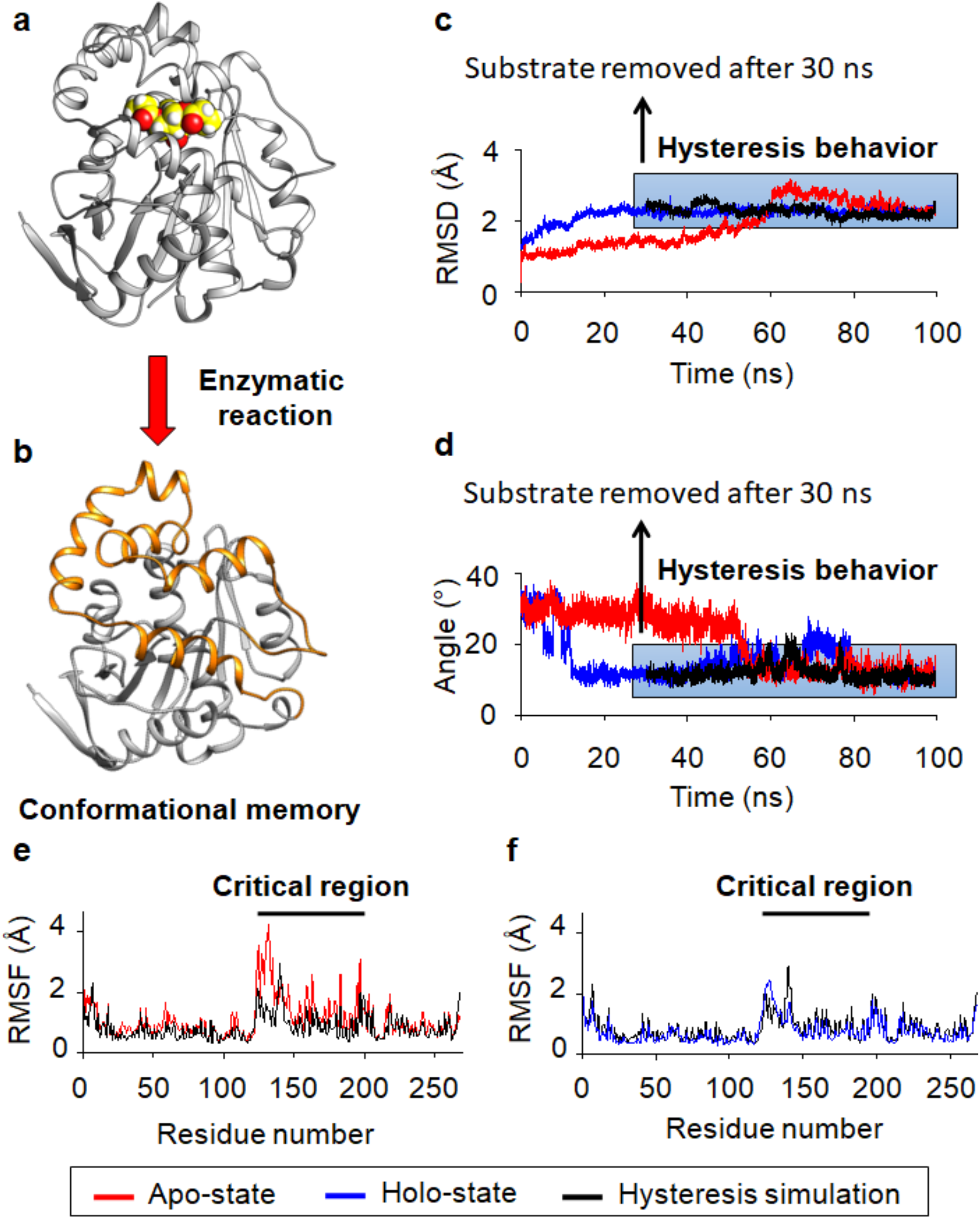
Hysteresis simulation of Ade1. To simulate a complete hydrolysis of the substrate, we removed it after 30 ns. In general, **(a)** Ade1 is kept open (E1 state, inactive catalytic conformation) for receiving tributyrin in its active site. **(b)** Next, tributyrin induces the closing of the cap domain (E2 state, active catalytic conformation) for advancing the substrate hydrolysis. Ade1 performs the complete hydrolysis of the tributyrin but it keeps the E2 state along with MD simulation. **(c)** Backbone root-mean-square deviation (RMSD) calculated for apo-state (red color), holo-state (blue color), and apo-state representing the hysteresis simulation (black color), which is characterized by structural maintenance of E2 state. **(d)** Closing angle (θ) was calculated using the coordinates formed by the main chain oxygen atoms of the V127, L27, and M194 residues. The closing angle plot was calculated for apo-state (red color), holo-state (blue color), and apo-state representing the hysteresis simulation (black color), which represents the closing angle of the E2 state. **(e)** Sidechain root-mean-square fluctuation (RMSF) was calculated for apo-state (red color) and apo-state representing the hysteresis simulation (black color). **(f)** Sidechain root-mean-square fluctuation (RMSF) was calculated for holo-state (blue color) and apo-state representing the hysteresis simulation (black color). Such results reveal that the hysteresis behavior is associated with the maintenance of active catalytic conformation (E2 state).

#### Ade1 S94 mutation to cysteine favors the E1 state (low activity conformation)

Some α/β-hydrolases that share structural similarities with Ade1 are cysteine hydrolases. Among them are the thioesterase GrgF (PDB ID 6LZH), the nonribosomal peptide synthetase ObiF1 (PDBID 6N8E), and the phosphotriesterase P91 (PDB ID 4ZI5). They are involved in the biosynthesis pathway of polyketide gregatin A ^74^, in antimicrobial obafluorin biosynthetic pathway^75^, and in hydrolysis of phosphotriester hydrolysis, respectively. Therefore, we produced an Ade1 mutant by changing the catalytic serine to a cysteine, Ade1_S94C_, for investigating its enzymatic activity. Interestingly, experimental assays demonstrated that Ade1_S94C_ lost completely its enzymatic activity (**Figures 3a, c** and **S8**). In order to enhance a molecular understanding for these results, we performed molecular dynamics simulation using a Ade1_S94C_ structure generated by changing the S94 to a cysteine in the wild type crystal structure. Since the structural coordinates of Ade1_S94C_ presented a distance ~3 Å between C94 and C118, we initially hypothesized the activity loss would be due to the formation of a disulfide bond. To test our hypothesis, we examined if, by restoring the nucleophilic nature of the thiol group of C94 in the presence of *β*-mercaptoethanol (reduction of disulfide bond C94-C118), we could detect any enzymatic activity. However, no activity was detected in our experimental set-up (data not shown).

To improve the molecular understanding of the inactivation caused by S94C site-directed mutation, we computed the angle formed by residues V127-L22-V194, sidechain RMSF, and backbone RMSD, and compared them to the wild type results (**Figure S16**). These results show that the structural flexibility of the cap domain of Ade1_S94C_ apo-state is impaired, as it exhibits a decrease of RMSF in relation to wild type (**Figure S16a**), which suggests that Ade1_S94C_ visits mainly one structural conformation during the MD simulation. According to the RMSD plot, Ade1_S94C_ is mainly in the E1 state (cap domain in the open conformation, **Figure S16b**). When compared with WT, Ade1_S94C_ exhibits an enzymatic cavity essentially kept in the open conformation with an angle of ~40° (**Figure S16c**). Interestingly, we observed that the Ade1_S94C_ holo-state also is affected in its RMSF (**Figure S16d**) and RMSD (**Figure S16e**) in relation to the Ade1 wild type. Sidechain RMSF, calculated using the Ade1_S94C_ holo-state, revealed that the cap domain flexibility is slightly bigger than the Ade1 wild type, in mean ~0.6 Å (**Figure S16d**). Furthermore, we observed that Ade1_S94C_ holo-sate is mainly in the E1 state and keeps the cap domain in an open conformation (low activity conformation) in contrast to the wild type that is mainly in the E2 state (high activity conformation) (**Figure S16e**).

Overall, we observed a negative effect under the mutant structure, which impaired the closing of the enzymatic catalytic cavity by the cap domain, a structural change necessary for advancing the substrate hydrolysis (**Figure S16f**) as seen for the cap domain of MenH ^72^. We also observed that Ade1_S94C_ delayed by 60 ns the closing of the cap domain when compared with WT, and it did not close completely to activate the catalytic triad (**Figure S16f**). In addition, we also noted an increase in the average distance between the mutant catalytic pair of 5.5 Å (C94 and H245) in comparison to the wild type of ~3 Å (S94 and H245), causing its functional inactivation (**Figure S16g** and **Figure S16h**). This result demonstrates that the proton transfer step is disrupted between C94 and H245. These results suggest that, even if it captures the substrate, the mutated enzyme is not able to close the cap domain for performing enzymatic catalysis progress.

## Conclusion

We described a novel esterase Ade1 from a metagenomic library of Amazon Dark Earth (ADE) soils from the Amazon Rainforest, in Brazil. Ade1 is a member of α/β-hydrolase superfamily with hydrolytic activity towards esteres with aliphatic group with less than 12 carbons and more than 3 carbons among the substrates tested in this work. Therefore, we can classify Ade1 as an esterase and not a lipase. Ade1 also hydrolases tributyrin (aliphatic chain with 3 carbons), tween-20 (aliphatic chain with 12 carbons) and *N*-hexanoyl-L-homoserine lactone (aliphatic chain with 6 carbons) showing a degree of substrate promiscuity. Some enzymes have been described to be moonlighting enzymes that have more than one biological function in the cell, depending on the moment and location of the enzyme ^76^. This class of enzymes are in general substrate promiscuous and multifunctional proteins ^76^, characteristics that Ade1 also shares. Despite the notorious difficulty to correlate biological function, enzymatic activity and identify specific substrates for members of the α/β-hydrolase superfamily based on three-dimensional structure, our biochemical assays and molecular dynamics simulations clearly demonstrate that Ade1 is a promiscuous esterase that has its function enhanced by the presence of cobalt. Ade1 enzymatic activity is unaffected by high concentrations of bivalent cations (concentrations ranging from 1 to 6 mM).

Interestingly, Ade1 has hysteresis behavior due to a presence of two conformational states, E1 and E2. Our MD simulations revealed that the E1 state has an open cap conformation (inactive catalytic conformation) while the E2 state has a closed cap domain (active catalytic conformation). Conformational equilibrium between E1 and E2, *K*_1_, is affected by the concentrations of the substrate and Co^2+^ ions that, in high concentrations, favor the E2 state. This model explains the characteristic transient burst of this enzyme, that leads to a sigmoidal curve in the kinetics data (V_o_ *versus* substrate concentration). This profile is not expected for monomeric enzymes that have one active site that bind one substrate and produce one product. Cobalt metal increases the Ade1 catalytic efficiency (*k*_cat_/ *K*_half_) probably by affecting the Ade1 conformation state equilibrium favoring the E2 state. This effect enhances the sigmoidal profile of the reaction curve that is observed in plots of the initial rate *versus* substrate. It is known that metabolic enzymes may have hysteresis behavior in order to regulate its enzymatic activity, which is dependent, in the case of Ade1, on the substrate and Co^2+^ concentrations ^69^.

In order to complement our experimental assays, we simulated Ade1 and Ade1_S94C_ using molecular dynamics simulation, which provided a clear insight to understand the molecular bases involving the mechanistic knowledge of enzymatic hysteresis. E1 is characterized by distancing between the catalytic pair S94/H245. However, E2 state is characterized by bringing the two catalytic residues closer, favoring proton transfer step, which is critical for the catalysis process. Interestingly, the mutation of the nucleophilic S94 to a cysteine, favoring the E1 state, abolished the catalytic activity. Based on that, we detailed the molecular bases of Ade1 as well as its functional and biochemical roles that probably is related to bacterial metabolism with hysteresis behavior. Moreover, this esterase may play a key role in an environment through its biological function probably in quorum-quenching activity and maintaining its enzymatic activity in high concentration of Co^2+^ (6mM).

Esterases are enzymes found in almost all living organisms, demonstrating their biological relevance and have been used as a tool in a wide range of applications in the food industry, pharmaceutical industries, and agriculture. Examples of this are the use of esterases in the degradation of industrial pollutants, plastics, and other toxic or xenobiotic compounds ^77,78^,^79^. Furthermore, they are also used in the synthesis of optically pure compounds, antioxidants, and perfumes ^80^. Understanding of the molecular bases, catalytic and structural mechanisms of Ade1 may be applied to other esterases of biotechnological, food, and/or pharmaceutical interest. In addition, our study enlarges the knowledge of the molecular mechanism of monomeric esterases with hysteresis behavior.

## Acknowledgement

Financial support was provided by the State of São Paulo Research Foundation (FAPESP, Fundação de Amparo à Pesquisa do Estado de São Paulo), Grant 2019/00195-2, 2020/04680-0. A. S. S. Acknowledges the Coordination for the Improvement of Higher Education Personnel (CAPES, Coordenação de Aperfeiçoamento de Pessoal de Nível Superior) for post-doctoral grant. T. C. V. Acknowledges the Coordination for the Improvement of Higher Education Personnel for institutional grant.

I would like to thank Professor Beny Spira and PhD Fernanda Nogales da Costa Vasconcelos for supplying *Chromobacterium violaceum* cells and helping with the quorum sensing assays.

