## supplemental information for "Molecular and functional basis of a novel Amazonian Dark Earth Esterase 1 (Ade1) with hysteresis behavior and quorum-quenching activity"

**Running title: Esterase with hysteretic behavior and quorum quenching activity**

To whom correspondence should be addressed: Prof. Cristiane Guzzo Carvalho, Department of Microbiology, Institute of Biomedical Science, University of São Paulo, Avenue Prof. Lineu Prestes, 1374, Cidade Universitária, 05508-900, São Paulo-SP, Brazil, Telephone: +55 11 3091-7298;

**Keywords:** metagenomic, X-ray crystallography, molecular dynamics simulations, bioinformatic, and enzymatic kinetics.

### Materials and Methods

#### Construction of soil metagenomic libraries and search for functional cloning to lipolytic activity

A metagenomic library from an Amazonian dark soil sample from Brazil was constructed in partnership with the Federal University of Amazonas, University of São Paulo and University of Brasília. Soil samples were collected, sieved, and stored at -20 °C. Total DNA was extracted and purified for generating a fosmid library using the pCC1FOS vector using Copy Control™ Fosmid Library Production commercial kit (Epicentre Biotechnologies, Chicago, Il, USA), with the vectors being transformed into *Escherichia coli* EPI300 (Epicentre). The DNA fragments cloned in the pCC1FOS vector were ~40,000 pb size.

In order to perform a functional selection, the clones were plated on Luria Bertani (LB) agar medium (1% tryptone, 0.5% yeast extract, 1% NaCl and 1.5% agar) supplemented with chloramphenicol 12.5 $\mu g.ml^{-1}$ and 1% tributyrin (1,3-bis-(butanoyloxy)-propane-2-yl-butanoate) as substrate (Sigma) emulsified with a sonicator and incubated at 37 ºC for 3 days. Lipolytic activity, hydrolysis of tributyrin, was verified by the presence of clear halos around the colonies (data not shown). In this first screening 14 out of 80,000 colonies tested presented hydrolyses halos. To confirm the observed phenotype the positive colonies were grown on LB medium containing arabinose (0,001%) to induce a high copy number of vectors according to OriV/TrfA amplification system [1]. One of them was chosen for further studies because it presented a higher hydrolysis halo (data not shown). In order to identify the gene coding the lipolytic activity sub-libraries constructions were performed using pUC18.

#### Sub-library construction of colonies that presented functional lipolytic activity

The plasmid was extracted from *Escherichia coli* EPI300 colony that presented lipolytic activity in the first screening, using the NucleoSpin Plasmid kit (Clontech Laboratories Inc., Mountain View, CA, USA). It was digested with *Hind*III for 2 h (Thermo Fisher Scientific Inc.) for generating DNA fragments around 2-10 kb, which were extracted and purified from agarose gels. DNA fragments were cloned into pUC18 cloning vectors (PubMed 6323249) and transformed into *E. coli* DH5α competent cells by electroporation. The transformed cells were plated on LB agar containing 100 μg.ml$^{-1}$ ampicillin, 200 μg.ml$^{-1}$ X-Gal (5-bromo-4-chloro-3-indolyl-β-D-galactopyranoside) and 100 μg.ml$^{-1}$ IPTG (isopropyl-β-D-thiogalactopyranoside) and incubated at 37 ºC overnight. White transformant colonies were screened by incubation in LB agar plates supplemented with 0.5% tributyrin emulsion and ampicillin for 72 h at 37 ºC. One out of 261 colonies had hydrolysis halo in the condition tested and was selected for a new sub-cloning procedure.

The plasmid was extracted from the *E. coli* DH5α that presented lipolytic activity using the NucleoSpin Plasmid kit (Clontech Laboratories Inc., Mountain View, CA, USA). Initially, the fragment cloned into the pUC18 vector was digested with the same restriction enzyme (*Hind*III) that confirmed a fragment size of ~8,000 pb. The same plasmid was digested with *Xba*I (Thermo Fisher Scientific Inc.) for generating DNA fragments ~1-8 kb, which were extracted and purified from agarose gels. DNA fragments were cloned into pUC18 cloning vectors (PubMed 6323249) using the same restriction enzyme and transformed into *E. coli* DH5α competent cells by electroporation. Transformed cells were plated on LB agar containing 100 μg.ml$^{-1}$ ampicillin, 200 μg.mL$^{-1}$ X-Gal (5-bromo-4-chloro-3-indolyl-β-D-galactopyranoside) and 100 μg.ml$^{-1}$ IPTG (isopropyl-β-D-thiogalactopyranoside) and incubated at 37 ºC overnight. White transformant colonies, a total of 332 colonies, were screened by incubation in LB agar plates supplemented with 0.5% tributyrin emulsion and ampicillin for 72 h at 37 ºC. Most of them had hydrolysis halo in the condition tested and 5 of them were randomly selected. The plasmid of the 5 colonies were extracted from the *E. coli* DH5α cells using the NucleoSpin Plasmid kit (Clontech Laboratories Inc., Mountain View, CA, USA). Initially, the fragments cloned into the pUC18 vector were digested with the same restriction enzyme (*Xba*I) that showed the same fragment size of ~2.3 kb for all plasmids extracted. It suggests that all of them have the same fragment and one was selected for further assays.

The DNA fragment cloned into the pUC18 was sequenced by the primer walking method using firstly M13-forward (5' CAGGAAACAGCTATGAC 3') and M13-reverse (5' CGCCAGGGTTTTCCCAGTCACGAC 3') and after two more oligonucleotides designed using the Gene Runner program [2]: Ade1-Foward (5´AGTTATCCACAGGAACACGG 3´) and Ade1-Reverse (5´ TACCTGGTCGCAGTTGCTG 3´). Sequencing reactions were performed at the Center for Research on the Human Genome and Stem Cells (Institute of Biosciences,

University of São Paulo) and analyzed using an ABI3730 DNA Analyzer (Applied Biosystems, Foster City, CA, USA). The DNA sequence was analysed and one Open Read Frame was identified that we named as *ade1* (Amazonian Dark Earths Esterase number 1).

#### *ade1* cloning into the expression vector pET28a

Gene encoding Ade1 was amplified by PCR using the following forward and reverse primers respectively: 5´-TTTTTCATATGCTATATGCTCAGGTCAACGGC-3´ (*Nde*I site underline) and 5´-TTTTGGATCCCTACACGCTTTGCCGGT-3´ (*Bam*HI site underline). PCR was performed using *Pfu* high fidelity DNA polymerase (Promega) in the following condition: an initial step of 2 min at 94 ºC, followed by two distinct cycles, firstly by 5 cycles of 94 ºC for 30 s, 51 ºC for 30 s and 68 ºC for 90 s, and secondly by 25 cycles of 94 ºC for 30 s, 58 ºC for 30 s and 68 ºC for 90 s. Final extension was 72 ºC for 10 min. PCR products were purified from the agarose gel and double-digested by *Nde*I and *Bam*HI restriction enzymes (ThermoFisher Scientific). Digested fragment was purified from the agarose gel and cloned into pET28a vector[3], previously digested with the same pair of enzymes (Novagen), resulting in the plasmid pET28a-Ade1. This construct generates a recombinant lipolytic enzyme with a $(His)_6$-tagged at the N-terminus of the protein for facilitating the protein purification step. Mixture ligation was inserted into *E. coli* DH5α by electroporation. Positive pET28a-Ade1 clones were confirmed by sequencing.

#### Ade1 classification by phylogenetic analysis

Amino acid sequence of Ade1 was subjected to protein phylogenetic analysis to classify it using the lipolytic data list constructed by Hitch and Clavel[4] They divided bacterial lipolytic enzymes by sequence similarities and function into thirty-five families[4]. Multiple sequence alignments (one per family and Ade1) were performed using the Clustal omega program[5] Multiple sequence alignment obtained were timing using Trimal -gt 0.5 for removing the sections that have up to 50% gaps. It was visually examined and edited in the Jalview 2.11.0 program. A phylogenetic tree was constructed with the neighbor-joining method using 1,000 bootstraps replicated by means of Molecular Evolutionary Genetics Analysis software (MEGA-X, version 5)[6] for determining the evolutive relations between Ade1 and bacterial lipolytic enzymes. All primary sequences, with exception Ade1, were obtained from UniProt database[7].

**Site-directed mutagenesis**

Ade1 was site-specifically mutated by PCR using the QuickChange Site-directed Mutagenesis Kit (Stratagene) using the primers below, which the mutation sites are highlighted in underline:

S94A-Fw: 5´ CCACGTCTTTGGCGTAGCAATGGGTGGGATGATCGCTCA 3´,

S94A-Rv: 5´ TGAGCGATCATCCCACCCATTGCTACGCCAAAGACGTGG 3´,

S94C-Fw: 5´ CCACGTCTTTGGCGTATGCATGGGTGGGATGATCGCTCA 3´,

S94C-Rv: 5´ TGAGCGATCATCCCACCCATGCATACGCCAAAGACGTGG 3´,

PCR products were digested with DnpI (Stratagene) for eliminating the parental methylated DNA and were introduced into *E. coli* XL1-Blue cells (Stratagene) by electroporation. Mutations sites were confirmed through DNA sequencing using T7 universal primers for pET28a.

**Expression and purification of Ade1 and Ade1 mutants**

Ade1, Ade1$_{S94A}$ and Ade1$_{S94C}$ were expressed in *E. coli* strain BL21(DE3)RP (Stratagene). Cells were grown at 37 °C in 2XTY medium (16 g.l$^{-1}$ bacto-tryptone, 10 g.l$^{-1}$ yeast extract and 5 g.l$^{-1}$ NaCl) supplemented with 50 µg.ml$^{-1}$ kanamycin (Gibco) and 30 µg.ml$^{-1}$ chloramphenicol (Sigma Aldrich). Expression of recombinant Ade1 was induced up to OD$_{600}$ 0.8, adding 1 mM isopropyl-β-D-1-thiogalactopyranoside (IPTG) and maintained in growth by 4 h. Bacterial cells were collected by centrifugation and resuspended in lysis buffer (50 mM Tris-HCl pH 7.5, 100 mM NaCl, 20% (w/v) sucrose, 10% glycerol (v/v), 0.03% (v/v) Triton X-100, and 0.03% (v/v) Tween 20) and then lysed by sonication. Ade1 was purified by affinity chromatography using a HisTrap Chelating HP column (GE Healthcare Life Sciences) previously equilibrated with Elution Buffer A (50 mM Tris-HCl pH 7.5, 100 mM NaCl, and 20 mM imidazole). Bound proteins were eluted in a range between 20 and 500 mM imidazole. Elution fractions were analyzed by 15% SDS-PAGE. All elution fractions containing Ade1 were concentrated through Amicon Ultra-4 Centrifugal filters (Merck Millipore) with a 3 kDa membrane cutoff. In order to remove the high concentrations of imidazole and salts, the protein solution was dialyzed in 10 mM Tris-HCl pH 7.5 and 10 mM NaCl.

**Lipolytic activity of Ade1 using Petri dishes**

Lipolytic activity (esterase or lipase) was tested using *E. coli* BL21(DE3)RP cells containing the expression vector for Ade1 (and $Ade1_{S94A}$ and $Ade1_{S94C}$). All assays were performed using 10 µl cell culture ($OD_{600}$ 0.8) on the top of LB solid medium supplemented with kanamycin 50 µg.ml$^{-1}$ (Gibco), chloramphenicol 30 µg.ml$^{-1}$ (Sigma Aldrich) and 0.5% (v/v) tributyrin (with a chain size of four carbons) or 0.5 % (v/v) triolein (with a chain size of eighteen carbons), in the presence of 1 mM IPTG. Tributyrin and triolein were emulsified in the medium by sonication. Petri dishes were incubated at 37 °C for 36 hours for observing the substrate hydrolysis.All assays were performed in triplicates.

**Tweenase activity**

Tweenase test was performed with an LB solid medium containing a final concentration of 4 mM $CaCl_2$, supplemented with 1% (v/v) Tween 20 or Tween 80. A volume of 10 µl (1 mg ml$^{-1}$) of each purified protein (Ade1 and $Ade1_{S94C}$) was applied on top of the solid medium and incubated at 37 °C for 24 hours to observe the precipitation of the reaction products (fatty acids) with the $Ca^{2+}$ in the medium, as described by Lee[8].

**Esterase kinetics assays by colorimetric measurements**

In order to determine the optimal enzymatic condition of Ade1, different substrates, such as *p*-nitrophenyl butyrate, *p*-nitrophenyl octanoate, *p*-nitrophenyl laurate and *p*-nitrophenyl palmitate were tested. They were dissolved in methanol and diluted ranging the concentrations from 20 to 150 nM. In order to verify the optimum pH of Ade1 for hydrolyzing the *p*-nitrophenyl octanoate, pH was ranged from 4.0 to 9.0. Furthermore, we monitored if the presence of different divalent metals ($Ca^{2+}$, $Co^{2+}$, $Mg^{2+}$, $Ni^{2+}$, and $Zi^{2+}$ at final concentrations of 0, 3 and 6 mM) could affect the enzymatic catalysis. All experiments were performed at different reaction buffers (100 mM Tris-HCl, 50 mM NaCl, 0.5% (v/v) Triton X-100, ranging the pH from 4.0 to 9.0) 600 µM *p*-nitrophenyl octanoate and Ade1. All enzymatic assays were performed in 96 well microplates at 30 °C. Kinetic reactions were started from the automated addition of Ade1, keeping as final condition was 100 mM Tris-HCl pH 8.0, 50 mM NaCl, 0.5% (v/v) Triton X-100, 600 µM *p*-nitrophenyl octanoate and 36 nM Ade1. Reaction product, *p*-nitrophenyl ester, was measured at absorbance 347 nm along 20 s for 5 min using a Synergy H1 Hybrid Multi-Mode Microplate Reader (BioTek, Winooski, VT,

USA). Calibration curves were performed using different concentrations of *p*-nitrophenyl ester. Measurement of absorbance was kept in the same conditions performed in the Ade1 kinetic assays. Linear regression and standard deviation calculations were performed for determining the initial reaction velocities through reaction production as time function. Michaelis-Menten and Hill-Langmuir models were compared for finding the best representation of the experimental data using the extra sum-of-squares F test in the Graph Path PRISM program[9]. All kinetic assays were performed in triplicates.

**Ade1 Crystallization, structure determination and refinement**

Initial crystallization trials were set up with the Crystal Screen 1, Crystal Screen 2, and Index reagent kits (Hampton Research) at 18 °C using the sitting-drop vapor-diffusion method in 24-well Linbro plates (Hampton Research). Each drop, consisting of 1.5 µL purified protein solution (15 mg.ml-1 into 10 mM Tris-HCL pH 7.5, 10 mM NaCl and 1 mM $MgCl_2$) and an equal volume of reservoir solution, was equilibrated against 300 µL reservoir solution. Crystals were obtained by the optimization of the initial crystallization conditions by changing the pH values and the precipitant concentration. The best crystallization condition, 0.1 Bis-Tris, pH 6.5, 0.1 M NaCl, 1.3 M $(NH_4)_2SO_4$, was added 4 mM tributyrin. Crystals were observed after 3 weeks.

All data were collected on a MicroMAX-007HR rotating-anode X-ray generator (Rigaku) equipped with an R-AXIS IV ++ detector, belonging to the Analytical Center, Chemistry Institute, USP. X-ray data were collected to 2.3 Å resolution at 100 K from a single crystal. The images were recorded with 7 min exposure using an oscillation range of 0.5°. The diffraction images were indexed, integrated, and scaled with iMOSFILM[10]. Initial phase was determined using PHASER [11] in CCP4i [12] by molecular replacement with a homology sequence deposited into to PDB database, PcaD (PDB ID 2XUA, resolution 1.9 Å) [13], which share identity and similarity of 29 and 44 %, respectively. An automated model was built using the ARP/wARp software [14] and the initial model was then refined through COOT [15] and REFMAC5 [16] packages. Alternative rounds of manual fitting were made from REFMAC5 to calculate the $R_{factor}$, $R_{free}$ and B-factor values. Three-dimensional structure of Ade1 was visualized using the graphic interface of COOT and PyMOL[17] and the final structure was validated using PROCHECK software [18].

***In vivo* activity of Ade1 as acyl-homoserine lactonase**

Acyl-homoserine lactonase (AHL) hydrolase *in vivo* activity was tested using *Chromobacterium violaceum* ATCC 12472. Purple-pigmented bacteria produce homoserine lactones (HSL), which mediate quorum-sensing and activate violacein production, a purple compound easily detected by spectrophotometry [19]. The amount of violacein in the culture medium was used as an indirect measure of HSL concentration. The assay was performed measuring the violacein production in the presence and absence of recombinant Ade1. The assay was performed using two different Ade1 concentrations (100 µg $ml^{-1}$ and 300 µg $ml^{-1}$) in 20 ml of LB culture medium containing *C. violaceum* with initial $OD_{600}$ of 0.1. Culture was incubated at 30 °C for 24 h at 150 rpm and aliquots of 100 µl were taken every 1 h. Each aliquot was centrifuged at 1,300 rpm for 4 min and the pellet was resuspended in 100 µl of DMSO to extract the total violacein. Afterward, new centrifugation was performed at 1,300 rpm for 4 min to eliminate the residual bacteria. The samples were then used to measure the absorbance at $OD_{585}$[19]. All assays were performed in triplicates.

**Determination of Ade1 molecular weight by Multi-angle light scattering coupled with size exclusion chromatography (SEC-MALS)**

Oligomeric state and molecular weight of Ade1 were determined through Multi-angle light scattering coupled with size exclusion chromatography (SEC-MALS). This assay was performed loading 500 µl of protein (2 $g.l^{-1}$) in the presence and absence of 6 mM $Co^{2+}$ into a Superdex 200 10/300 GL column (GE Healthcare) pre-equilibrated in 50 mM Tris-HCl pH 8.0 and 400 mM NaCl and the column was coupled to miniDAWN TREOS multi-angles light scattering detector/multi-angle light scattering detector (MALS) and Optilab TrRX refractive index detector/refractive index detector. All reactions were kept on ice for 1 h before being applied to the size exclusion column. The reactions were eluted at 0.5 $ml^{-1}$.min at 23 °C. Data was analyzed using OriginLab software. The assays were performed in triplicate.

**System setup for molecular dynamics simulations**

Next, we ran molecular dynamics simulations to study the structural aspects of wild-type and $Ade1_{S94C}$. Ade1 structure (PDB ID 6EB3) was prepared using the UCSF chimera tool[20] by removing co-crystallized hetero groups and water molecules.

$Ade1_{S94C}$ was obtained from a site-directed mutation in wild-type Ade1 using the Maestro software (academic v. 2020-1) [21]. From the Ade1 structure, protonation states of ionizable residues were computed in an aqueous implicitly environment at pH 8.0 from the Maestro software academic v. 2020-1 [21] using PROPKA module [22]. Then, all glutamic and aspartic residues were represented as unprotonated; H12, H47, H106, H125, H163, H165, H245 were designed as a δ-tautomer; H89, H172, and H189 were modeled as an ε-tautomer; all glutamic residues were kept with neutral charge; arginine and lysine residues were assumed with a positive charge; the N- and C-terminal, corresponding to M1 and V268, were converted to charged groups. To build a holo-state of Ade1 and $Ade1_{S94C}$, we docked the substrate into the binding site using as reference the crystal reaction product. To that end, we obtained the three-structure representation of the substrate tributyrin employing the conjugate gradient algorithm associated with the Merck molecular force field (MMFF94s) [23] using Avogadro software v. 1.2.1 [24]. Subsequently, the substrate was submitted to a genetic algorithm (GA) for performing the molecular docking into binding pockets of Ade1 and $Ade1_{S94C}$ using the genetic optimization for ligand docking software (GOLD, v. 2019-2) [25]. The poses were evaluated by a force field-based fitness function (gold score) and selected for molecular dynamics studies.

All molecular dynamics runs were performed in the Groningen machine for chemical simulation software (GROMACS, v. 5.1.5) [26,27], using optimized potentials for liquid simulations for all atoms (OPLS-AA) force field [28]. Substrate topology was built in LigParGen web-based service [29] and the 1.14*CM1A charges[30] were kept on substrate atoms. All systems were then explicitly solvated with TIP3P water models in a cubic box and neutralized with the addition of 5 sodium ions and minimized until to reach a maximum force of 10.0 $kJ.mol^{-1}$ or a maximum number of steps in 5000. Final dimensions were approximately 72.6 × 72.6 × 72.6 $Å^3$, including 5 sodium ions and around 11000 *TIP3P* waters. The systems were equilibrated consecutively in isothermal-isochoric (NVT) and isothermal-isobaric (1 bar; NpT) ensembles, both at temperature 300 K for 1 ns. All simulations were then performed in a periodic cubic box considering the minimum distance of 1.0 nm between any protein atom and cubic box walls. Molecular dynamics runs were performed for 100 ns. Finally, to determine the closing angle, we used the coordinates formed by the main chain oxygen atoms from the V127, L27, and M194 residues. Distance calculation, hydrogen bonding occupancy percentage, root-mean-square deviation (RMSD) and root-mean-square fluctuation (RMSF) were determined using the GROMACS modules and virtual molecular dynamics (VMD; v. 1.9.1) [31].

***Accession Number***

The nucleotide sequence obtained in this study has been deposited in the GenBank database under the accession number MW341220. The structure of Ade1 protein was deposited in the PDB databank under code 6EB3.

### Figures

```
Ade1   1    MLYAQVNGINLHYEIEGQGQPLLLIMGLGAPAAAWDPIFVQTLTKTHQVIIYDNRGTGLS  60
            M YA +NGI LHYE EG G PLL + GLG PA AWDP  VQ +   +QVI YDNRGTGLS
Sbjct  1    MSYAHINGIRLHYETEGHGPPLLFVAGLGQPAVAWDPALVQQMATQYQVITYDNRGTGLS  60

Ade1   61   DKPDMPYSIAMFASDAVGLLDALNIPRAHVFGVSMGGMIAQELAIHYPQRVASLILGCTT  120
            DKPD PY+IA+FASDAVGLLD LNIPRAHVFGVSMGGMIAQEL I+   RVASL LGCTT
```

```
Sbjct  61    DKPDEPYTIALFASDAVGLLDTLNIPRAHVFGVSMGGMIAQELGINAASRVASLTLGCTT  120

Ade1   121   PGGKHAVPAPPESLKALEGRAGLTPEEAIREGWKLSFSEEFIHTHKAELEAHIPRLLAQL  180
             PGG++AV APPESLK LEGRAG+TPE A R+GWKLSFS++FI TH+AELE H+ R L Q+
Sbjct  121   PGGRNAVQAPPESLKMLEGRAGMTPEAAARDGWKLSFSDDFIRTHQAELEGHMRRGLTQV  180

Ade1   181   TPRFAYERHFQATMTLRVFKQLKEIQAPTLVATGRDDMLIPAVNSEILAREIPGAELAIF  240
             TPRFAYERHFQAT+TLRVFKQLKEI APTLV TG+DD+LIPA NSEILAREIPGAEL  +
Sbjct  181   TPRFAYERHFQATLTLRVFKQLKEITAPTLVITGKDDILIPAANSEILAREIPGAELTLL  240

Ade1   241   ESAGHGFVTSAREPFLKVLKEFLARQ  266
             ++AGHGF  SARE F+ V +EFL R
Sbjct  241   DNAGHGFFISARERFVPVFQEFLTRH  266
```

**Figure S1. Primary sequence alignment of Ade1 and HOP18_27540 (GenBank: NOT58366.1).** Sequence alignment was performed using BLAST (Basic Local Alignment Search Tool) of NCBI (National Center for Biotechnology Information). Ade1 and HOP18_27540 share identity and similarity of 74 and 84%, respectively, with an e-value of $2.10^{-143}$.

```
PcaD   1    MPYAAVNGTELHYRIDGERHGNAPWIVLSNSLGTDLSMWAP-QVAALSKHFRVLRYDTRG  59
            M YA VNG  LHY I+G+       ++L   LG   + W P  V  L+K  +V+ YD RG
Ade1   1    MLYAQVNGINLHYEIEGQGQP----LLLIMGLGAPAAAWDPIFVQTLTKTHQVIIYDNRG  56

PcaD   60   HGHSEAPKGPYTIEQLTGDVLGLMDTLKIARANFCGLSMGGLTGVALAARHADRIERVAL  119
             G S+ P  PY+I     D +GL+D L I RA+  G+SMGG+    LA  +  R+  + L
Ade1   57   TGLSDKPDMPYSIAMFASDAVGLLDALNIPRAHVFGVSMGGMIAQELAIHYPQRVASLIL  116

PcaD   120  -CNTAARIGSPEVWVPRAVKARTEGMHALA--DAVLPRW---FTADYMEREPVVLA--MI  171
             C T    G   V  P       EG   L   +A+   W   F+ +++      L   +
Ade1   117  GCTTPG--GKHAVPAPPESLKALEGRAGLTPEEAIREGWKLSFSEEFIHTHKAELEAHIP  174

PcaD   172  RDVFVHTDKEGYASNCEAIDAADLRPEAPGIKVPALVISGTHDLAATPAQGRELAQAIAG  231
            R +   T +  Y  + +A     +  +   I+ P LV +G  D+         LA+ I G
Ade1   175  RLLAQLTPRFAYERHFQATMTLRVFKQLKEIQAPTLVATGRDDMLIPAVNSEILAREIPG  234

PcaD   232  ARYVELD-ASHISNIERADAFTKTVVDFLTEQ  262
            A     + A H   +  A  F K + +FL  Q
Ade1   235  AELAIFESAGHGFVTSAREPFLKVLKEFLARQ  266
```

**Figure S2. Primary sequence alignment of Ade1 and PcaD (UniProtKB - Q13KT2 (Q13KT2_PARXL).** Both Ade1 and PcaD share 29 and 44 % of identity and similarity, respectively, and an e-value of $6.10^{-28}$.

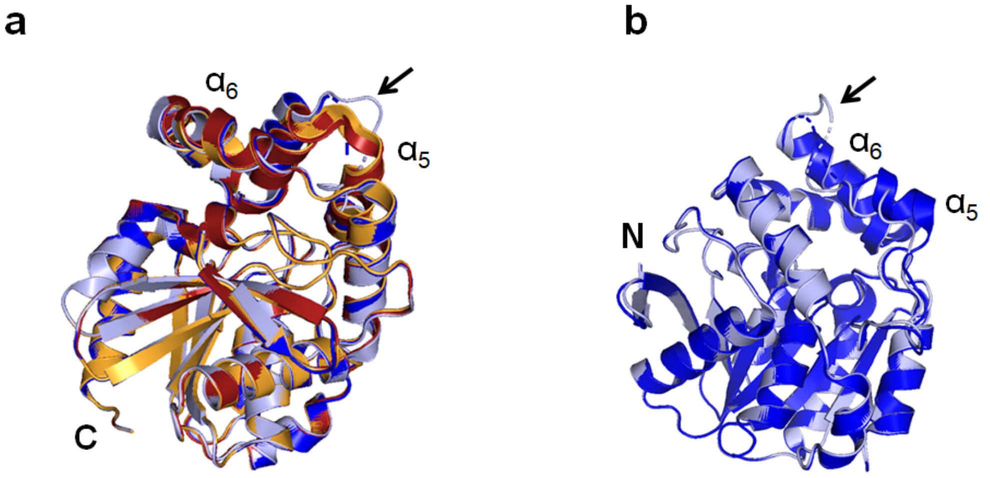


**Figure S3. Structural superposition of the four Ade1 chains (A-D) into the asymmetric unit (ASU). (a)** Superposition of the four Ade1 structures contained into the ASU. Chains A, B, C, and D are colored in light blue, red, dark blue and orange colors, respectively. **(b)** superposition of chains A and D of Ade1. Both panels show that, between $\alpha_5$ and $\alpha_6$, there is an unstructured region localized in the residues 138-140 (chain A) and 138-143 (chain C).

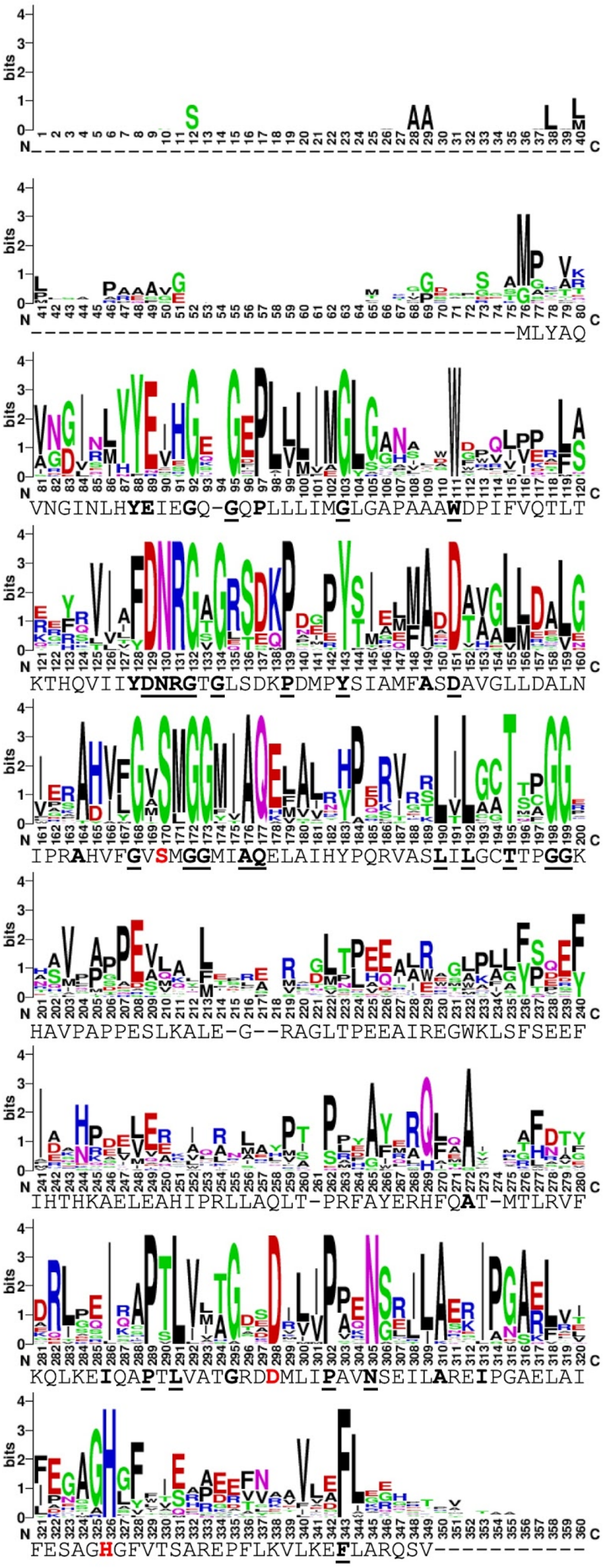

**Figure S4. Multiple sequence alignment representation of Ade1 analyzed in WEBLOGO web-based service.** Multiple sequence alignment was performed with proteins that share identity more than 45% and cover above 80% relative to Ade1

sequence to make a WEBLOGO presentation. Residues in bold are highly conserved and those underlined are absolutely conserved. Catalytic triad are colored in red, S94, D217 and H245.

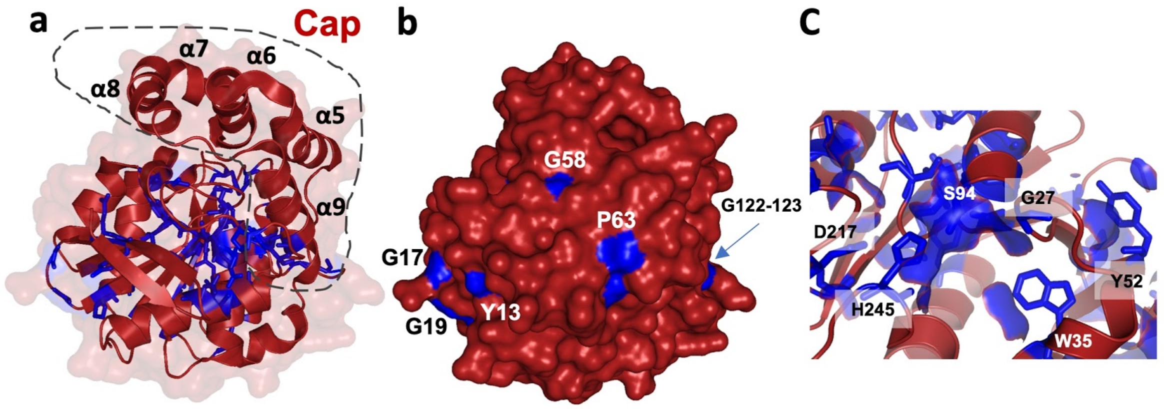


**Figure S5. Ade1 conserved residues mapped on the 3D-structure. (a)** Conserved residues within Ade1 orthologues are colored in blue and are described in details in **Figure S4**. Most of them are located at the Ade1 core domain with exception of G122, G123, and A192 residues that are located in the cap domain. **(b)** Conserved residues in the Ade1 structure that are exposed to solvent are colored in blue. **(c)** Conserved residues located in the catalytic pocket showing that S94 are exposed to the solvent.

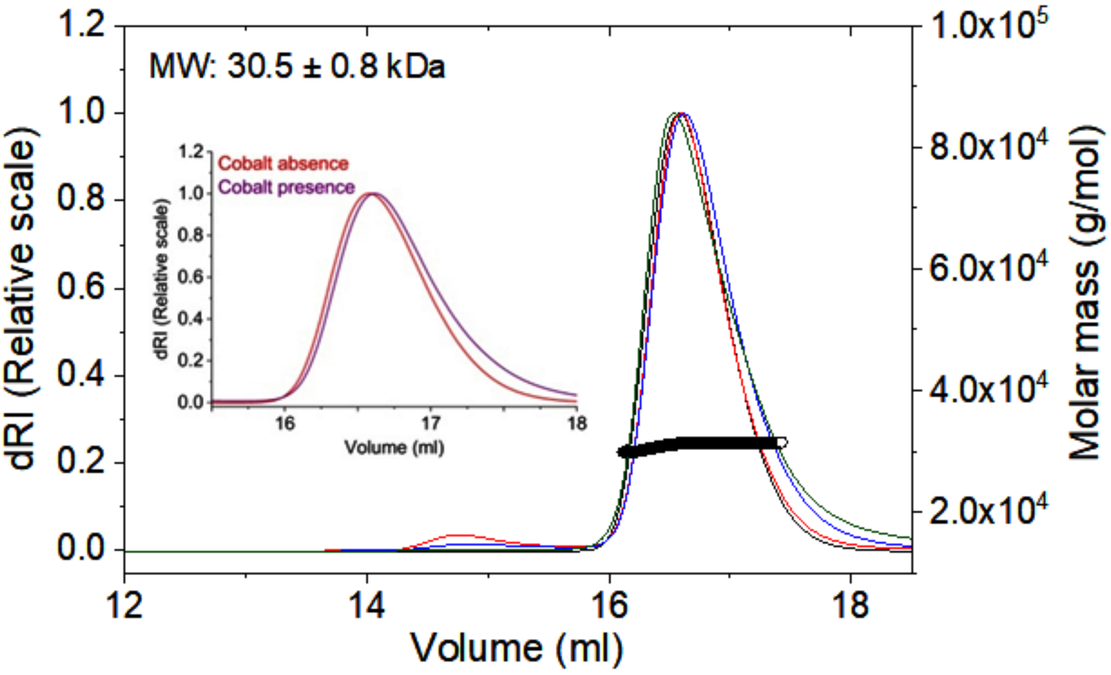


**Figure S6. Molecular weight of Ade1 determined by molecular exchange chromatography coupled to a multi-angle light scattering detector (SEC-MALS).** Determination of oligomeric states and molecular mass of Ade1 by molecular exchange chromatography coupled to a multi-angle light scattering detector (SEC-MALS). In this experiment was applied 550 µl of sample at 2 $g.l^{-1}$ Ade1 (Tris-HCl pH 8 50 mM and 400 mM NaCl, in absence and presence of 6 mM $Co^{2+}$). Big graphic shows the profile of all runs (Ade1 in absence and presence of $Co^{2+}$). Only one pick of Ade1 (monomer) was observed on the chromatogram, which revealed that Ade1 has a molecular weight of 30.5 ± 0.8 kDa. Little graphic shows Ade1 in absence and presence of $Co^{2+}$ (red and purple colors). All runs were performed in triplicate.

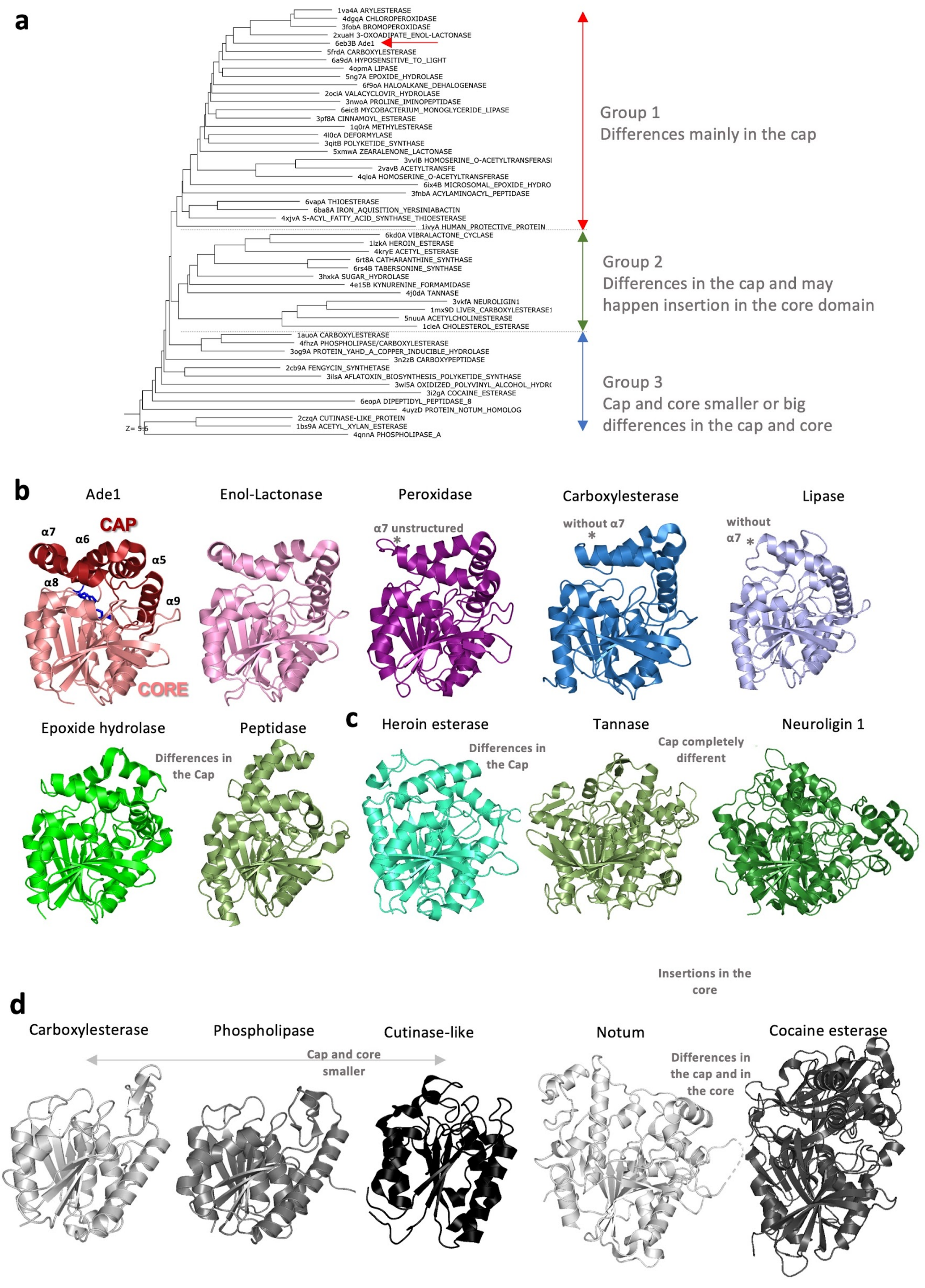

a
1va4A ARYLESTERASE
4dgqA CHLOROPEROXIDASE
3fobA BROMOPEROXIDASE
2xuaH 3-OXOADIPATE_ENOL-LACTONASE
6eb3B Ade1
5frdA CARBOXYLESTERASE
6a9dA HYPOSENSITIVE_TO_LIGHT
4opmA LIPASE
5ng7A EPOXIDE_HYDROLASE
6f9oA HALOALKANE_DEHALOGENASE
2ociA VALACYCLOVIR_HYDROLASE
3nwoA PROLINE_IMINOPEPTIDASE
6eicB MYCOBACTERIUM_MONOGLYCERIDE_LIPASE
3pf8A CINNAMOYL_ESTERASE
1q0rA METHYLESTERASE
4l0cA DEFORMYLASE
3qitB POLYKETIDE_SYNTHASE
5xmwA ZEARALENONE_LACTONASE
3vvlB HOMOSERINE_O-ACETYLTRANSFERASI
2vavB ACETYLTRANSFE
4qloA HOMOSERINE_O-ACETYLTRANSFERASE
6ix4B MICROSOMAL_EPOXIDE_HYDRO
3fnbA ACYLAMINOACYL_PEPTIDASE
6vapA THIOESTERASE
6ba8A IRON_AQUISITION_YERSINIABACTIN
4xjvA S-ACYL_FATTY_ACID_SYNTHASE_THIOESTERASE
1ivyA HUMAN_PROTECTIVE_PROTEIN
6kd0A VIBRALACTONE_CYCLASE
1lzkA HEROIN_ESTERASE
4kryE ACETYL_ESTERASE
6rt8A CATHARANTHINE_SYNTHASE
6rs4B TABERSONINE_SYNTHASE
3hxkA SUGAR_HYDROLASE
4e15B KYNURENINE_FORMAMIDASE
4j0dA TANNASE
3vkfA NEUROLIGIN1
1mx9D LIVER_CARBOXYLESTERASE1
5nuuA ACETYLCHOLINESTERASE
1cleA CHOLESTEROL_ESTERASE
1auoA CARBOXYLESTERASE
4fhzA PHOSPHOLIPASE/CARBOXYLESTERASE
3og9A PROTEIN_YAHD_A_COPPER_INDUCIBLE_HYDROLASE
3n2zB CARBOXYPEPTIDASE
2cb9A FENGYCIN_SYNTHETASE
3ilsA AFLATOXIN_BIOSYNTHESIS_POLYKETIDE_SYNTHASE
3wl5A OXIDIZED_POLYVINYL_ALCOHOL_HYDR(
3i2gA COCAINE_ESTERASE
6eopA DIPEPTIDYL_PEPTIDASE_8
4uyzD PROTEIN_NOTUM_HOMOLOG
2czqA CUTINASE-LIKE_PROTEIN
1bs9A ACETYL_XYLAN_ESTERASE
4qnnA PHOSPHOLIPASE_A
Z= 5.6
Group 1
Differences mainly in the cap
Group 2
Differences in the cap and may happen insertion in the core domain
Group 3
Cap and core smaller or big differences in the cap and core
b
Ade1
Enol-Lactonase
Peroxidase
Carboxylesterase
Lipase
CAP
α7
α6
α5
α8
α9
CORE
α7 unstructured
without α7
without α7
Epoxide hydrolase
Peptidase
Differences in the Cap
c
Heroin esterase
Tannase
Neuroligin 1
Differences in the Cap
Cap completely different
Insertions in the core
d
Carboxylesterase
Phospholipase
Cutinase-like
Notum
Cocaine esterase
Cap and core smaller
Differences in the cap and in the core

**Figure S7. Structural comparison of the cap domain between α/β hydrolases proteins. (a)** Structural similarity dendrogram using structures similar to Ade1. The dendrogram is derived by average linkage clustering of the structural similarity matrix (Dali Z-scores). It shows three groups delimited by the double arrow (red, green, and blue, respectively). Group 1 is characterized by slight differences only in the cap domain). Group 2 has differences in the cap and may happen insertion in the core domain. Group 3 shows two sub-groups presenting smaller or bigger differences in the cap and core domains. Red arrow indicates Ade1. The painels **(b), (c), and (d)** show the crystal structures of different members of α/β-hydrolases representants of the group 1, 2 and 3, respectively. **(b)** Ade1 (PDB ID 6EB3, chain B), enol-lactonase (PDB ID 2XUA, chain H), peroxidase (PDB ID 4DGQ, chain A), carboxylesterase (PDB ID 5FRD, chain A), lipase (PDB ID 4OPM, chain A), epoxide hydrolase (PDB ID 5NG7, chain A), proline peptidase (PDB ID 3NWO, chain A); **(c)** heroin esterase (PDB ID 1LZK, chain A), tannase (PDB ID 4J0D, chain A), neuroligin-1 (PDB ID 3VKF, chain A); **(d)** carboxylesterase (PDB ID 1AU0, chain A), phospholipase (PDB ID 4FHZ, chain A), cutinase-like protein (PDB ID 2CZQ, chain A), notum (PDB ID 4UYZ, chain D) and cocaine esterase (PDB ID 3I2G, chain A).

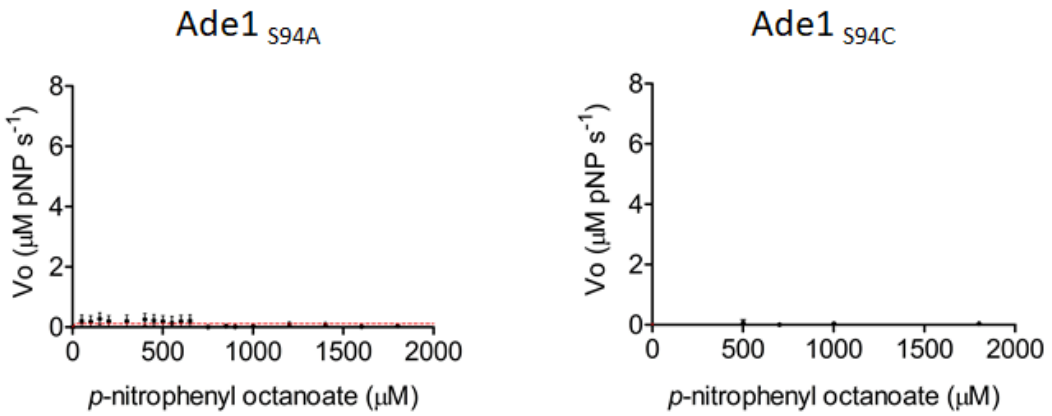


**Figure S8. Ade1 mutants (S94A and S94C) kinetic activity assays**. $Ade1_{S94A}$ and $Ade1_{S94C}$ mutants of Ade1 did not present enzymatic catalysis using *p*-nitrophenyl octanoate as substrate.

**Figure S9. Ade1 concentration as a function of enzymatic activity**. Reactions were performed at 30 °C in 100 mM Tris-HCl pH 8.0, 50 mM NaCl, 0.5% (v/v) Triton X-100, 600 µM *p*-NP C8, ranging the Ade1 concentration between 20 nM and 80 nM. Calibration curves were performed using different concentrations of *p*-nitrophenyl ester and the measurement of absorbance at 347 nm in the same buffer conditions used to Ade1 kinetic assays. Each assay was performed in triplicate and the average

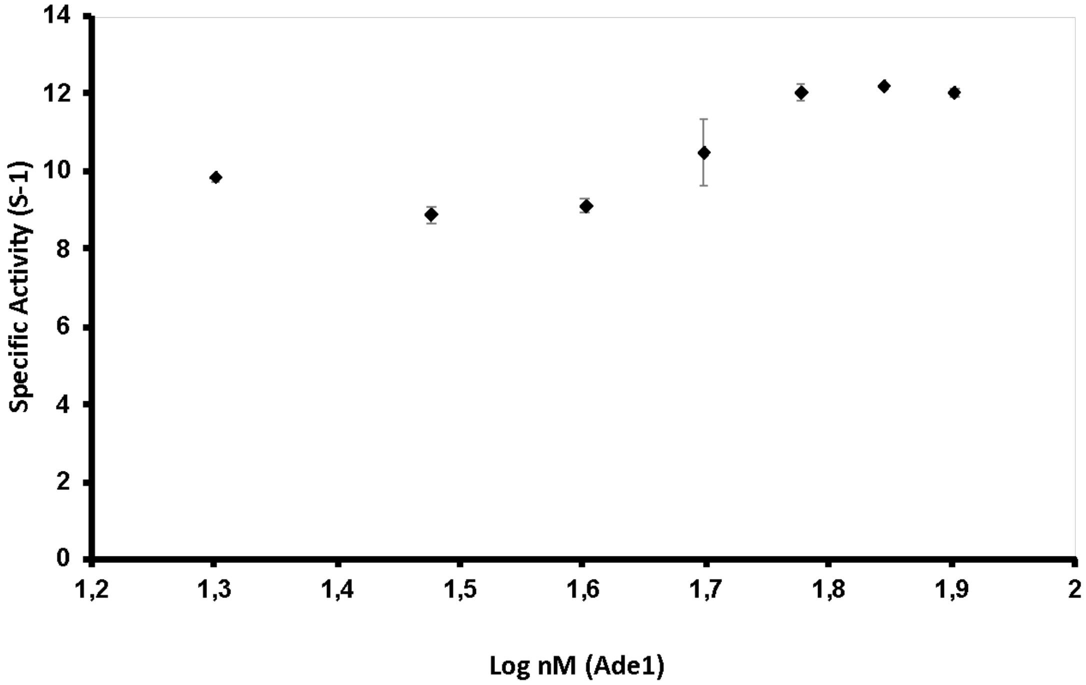


values were used for constructing the kinetic curve and determined the kinetic curve, standard deviations and enzymatic parameters.

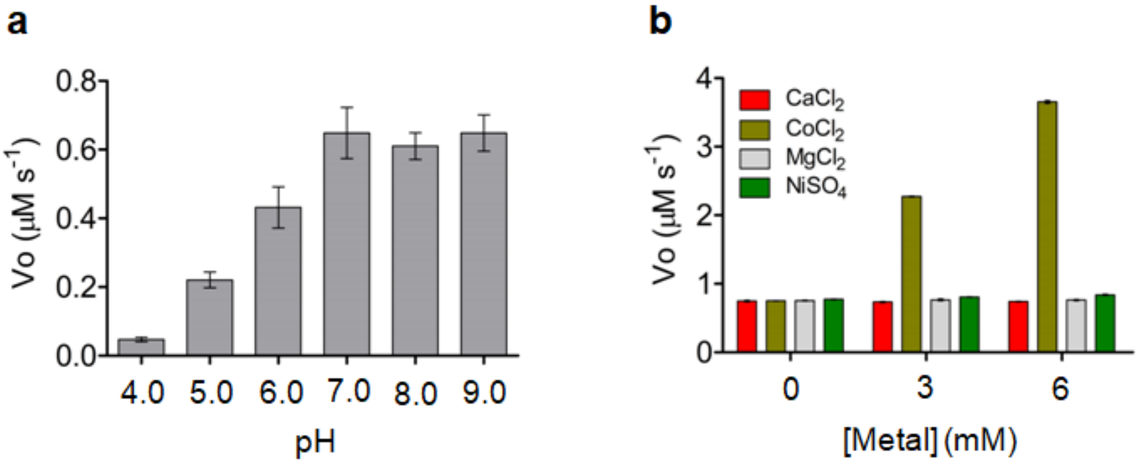


**Figure S10. Biochemical assays of Ade1.** Initial hydrolysis velocity of 600 μM *p*-nitrophenyl octanoate at different **(a)** pH (4.0 to 9.0) and **(b)** metals ($Ca^{2+}$, $Co^{2+}$, $Mg^{2+}$, and $Ni^{2+}$). In panel **(b)**, Ade1 was previously dialyzed at 1 mM EDTA.

**a**

**b**

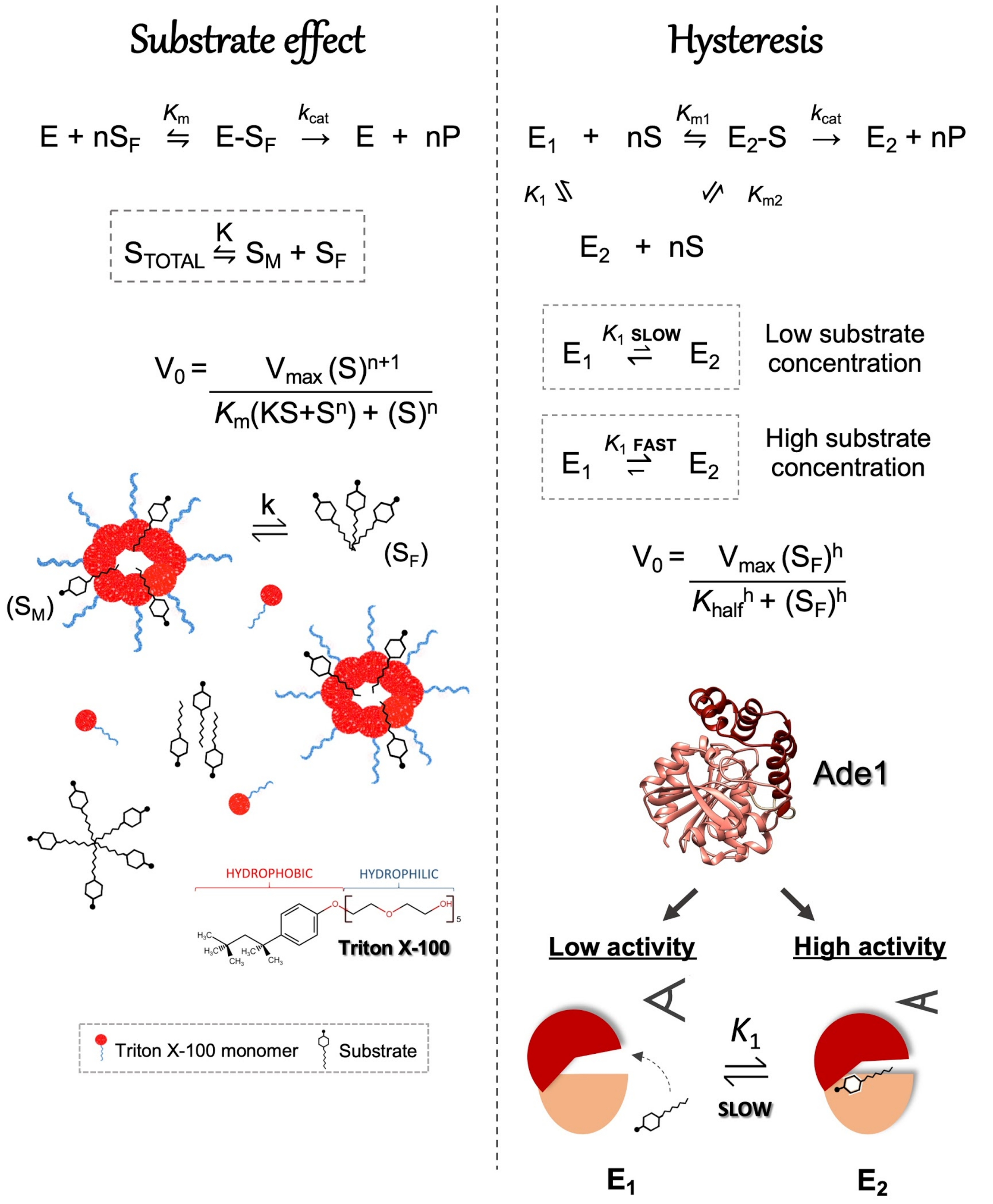


**Figure S11. Hypotheses that could explain the sigmoidal curve observed in the initial rate *versus* [S] plots. (a)** Substrate effect and **(b)** Hysteresis. Panel **(a)** shows substrate in two states equilibrium: $S_M$ (docked substrate into the Triton micelle) and $S_F$ (substrate outside the micelle, also known as free substrate. It could be grouped into two or more monomers). Thus, Ade1 cleavages only the free substrate and consequently shows a non-hyperbolic curve of cooperativity (sigmoid). $K_1$ is defined as a dissociation constant of the substrate inside and outside of the Triton X-100 micelles. Panel **(b)** shows the enzyme in two conformational states equilibrium, E1 (inactive catalytic state) and E2 (active catalytic state). Ade1 E1 state shows the cap domain in an open conformation, whereas E2 state is characterized by closing the cap domain. Substrate affects the equilibrium between both states favoring the E2 state (more details, **Figures S13** and **S14**).

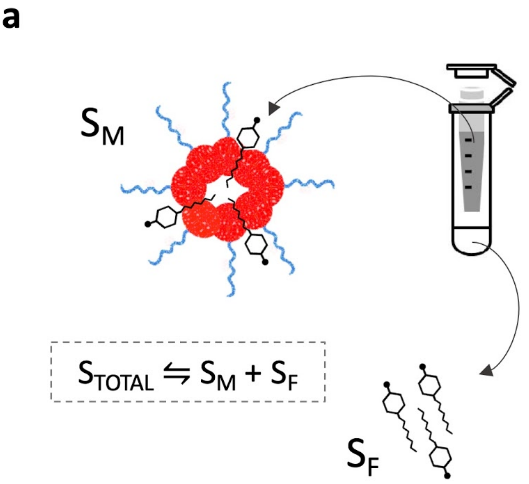


c

| *p*-nitrophenyl octanoate (µM) | |
|---|---|
| TOTAL | FREE |
| 700 | 255.71 |
| 750 | 273.975 |
| 800 | 292.24 |
| 850 | 310.505 |
| 900 | 328.77 |
| 1000 | 365.3 |
| 1200 | 438.36 |
| 1400 | 511.42 |
| 1600 | 584.48 |
| 1800 | 657.54 |
| 2000 | 730.6 |

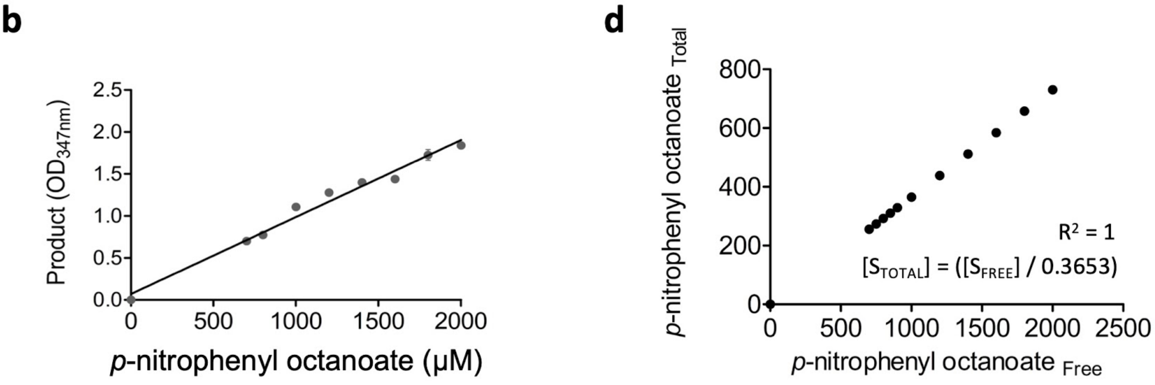


**Figure S12. Detection of free *p*-nitrophenyl octanoate in the mix enzymatic reaction.** (**a**) The *p*-nitrophenyl octanoate is in two states, $S_M$ (docked substrate into the Triton micelle) and $S_F$ (substrate outside the micelle, also known as free substrate. It could be grouped into two or more monomers). To determine the free *p*-nitrophenyl octanoate concentration in the reaction mix we incubated different concentration of *p*-nitrophenyl octanoate in the same reaction buffer for 1 hour at 25 °C, then it was centrifuged in Amicon Ultra-4 Centrifugal filters (Merck Millipore with 3 kDa of cutoff) to remove the Triton X-100 micelles. (**b**) The $S_F$ concentrations in the filtrated were determined by their total hydrolysis using Ade1. The product (*p*-nitrophenyl) was measured (OD 347 nm). A calibration curve was then used to calculate the concentration of *p*-nitrophenyl in relation to OD 347 nm. (**c**) Shows the total substrate used in each reaction (TOTAL) and the free substrate concentration (FREE). (**d**) The total substrate concentration and free substrate concentration are linear correlated.

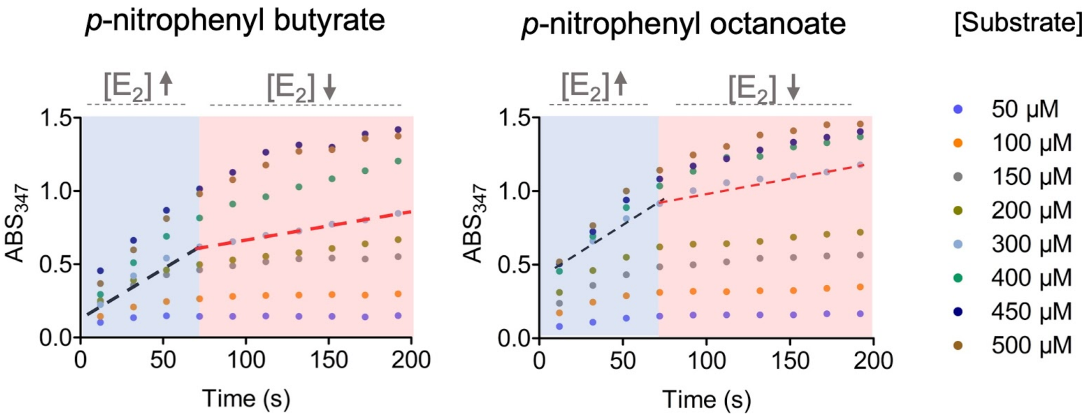


**Figure S13. Ade1 kinetic activity.** Ade1 shows hydrolysis of *p*-nitrophenyl butyrate (left) and *p*-nitrophenyl octanoate (right). According to both graphics, Ade1 has higher initial reaction velocity (black dotted lines) than steady-state velocity (red dotted lines). They are directly associated with high and low reaction velocities, respectively. Thus, Ade1 is classified as transient burst. This effect may be due to the concentration of the E2 state ([E2]) that is higher in high concentration of substrate and lower in low substrate concentration. This fluctuation of [E1] and [E2] concentrations in the reaction course is probably due to the effect of the substrate in the equilibrium constant ($K_1$, shown in **Figure S11**).

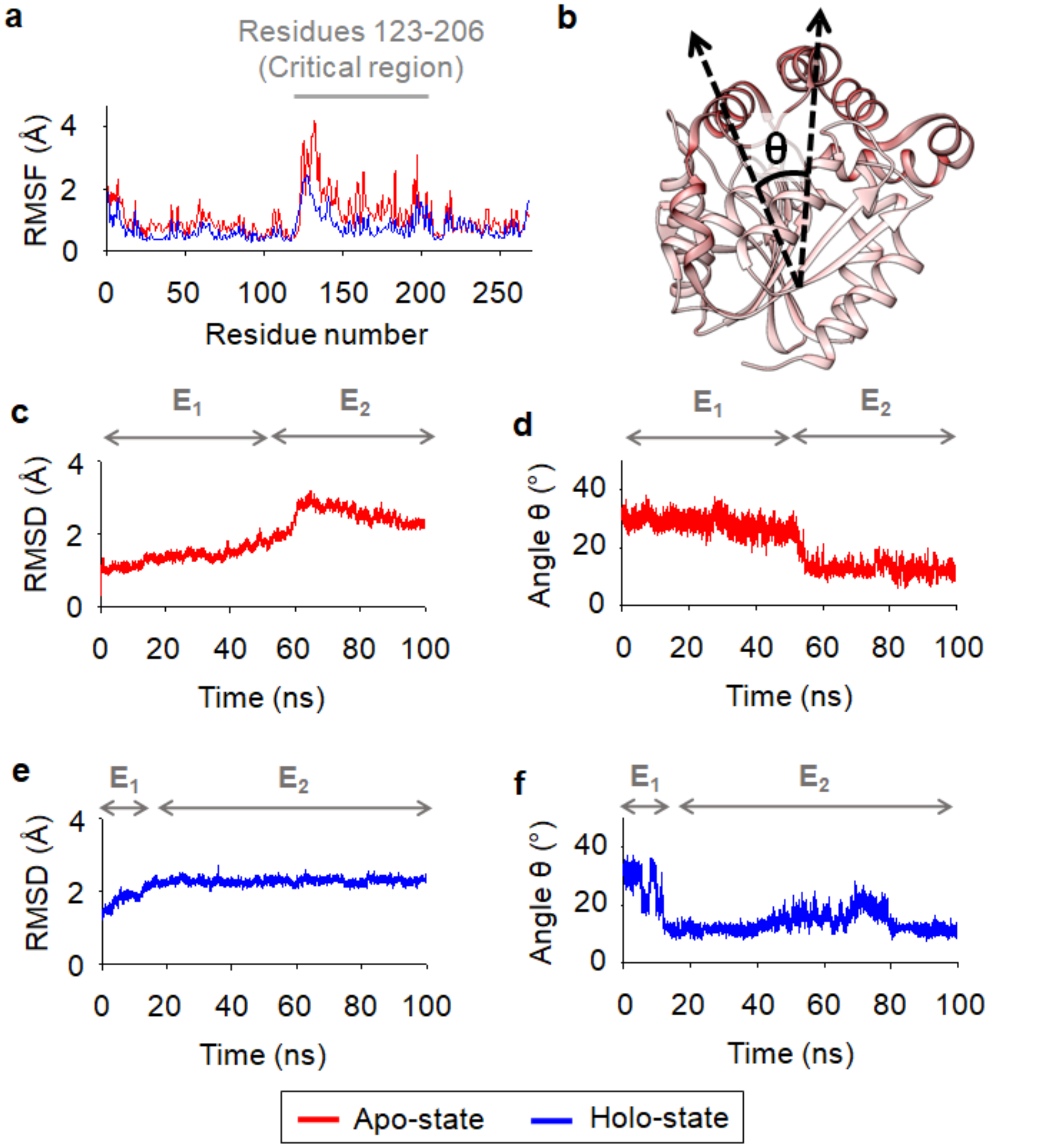


**Figure S14. Substrate induces the cap domain closing for the catalysis pathway advance.** The opening and closing of the cap domain are represented by the E1 and E2 states, respectively. **(a)** Esterase per-residue root-mean-square fluctuation (RMSF) plot according to color (red, apo-state; blue, holo-state). **(b)** The high fluctuation region, also called the critical region, is represented by residues 123-206. This region is highlighted in a dark red color. The closing angle (θ) was calculated using the coordinates formed by the main chain oxygen atoms of the V127, L27, and M194 residues. Closing angle plot as time function is shown on panel **(d)** and **(f)**. **(c)** Backbone root-mean-square deviation (RMSD) plot of apo-state. **(d)** Cap domain closing dynamics of Ade1 apo-state. **(e)** Backbone root-mean-square deviation (RMSD) plot of holo-state. **(f)** Cap domain closing dynamics of Ade1 holo-state.

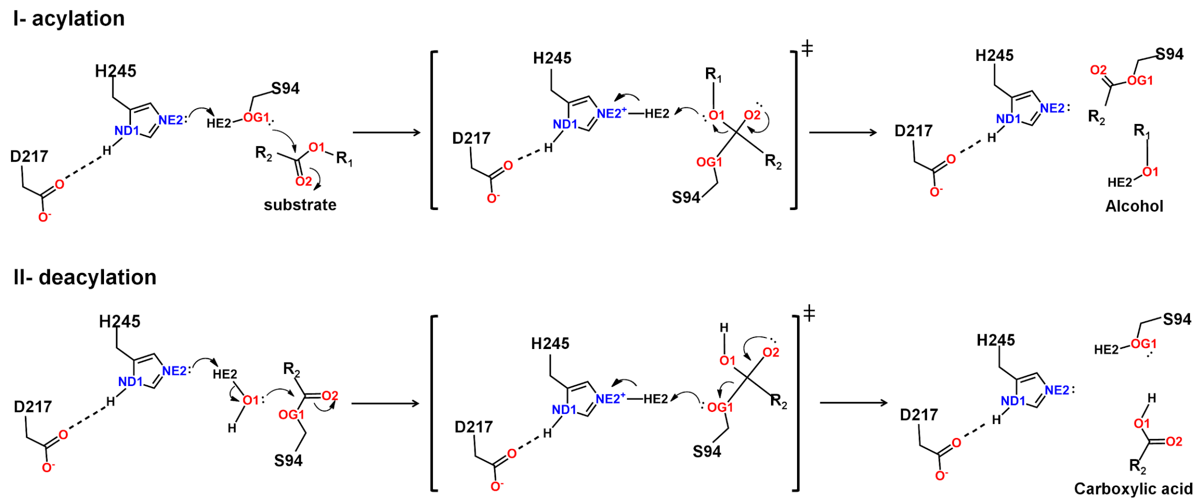


**Figure S15. Proposed enzymatic catalysis.** Reaction mechanism is characterized by theI-acylation and II-deacylation stages. In the acylation stage, S94 transfers a proton to H245, forming an ionic pair ($S94^-$/$H245^+$). Next, the charged serine attacks the carbonyl carbon atom of the substrate to produce a transient tetrahedral intermediate (TTI). The TTI oxygen electron pair forms a π bond and the H245 residue can donate a proton Hε2 to TTI, forming the reaction product $R_1$-OH. In the deacylation stage, the H245 acts as a Lewis base activating a water molecule. It attacks the carbonyl carbon forming a transient state. An oxygen electron pair forms a π bond and the initial state of the catalytic serine is restored and the product reaction $R_2$–COOH is formed.

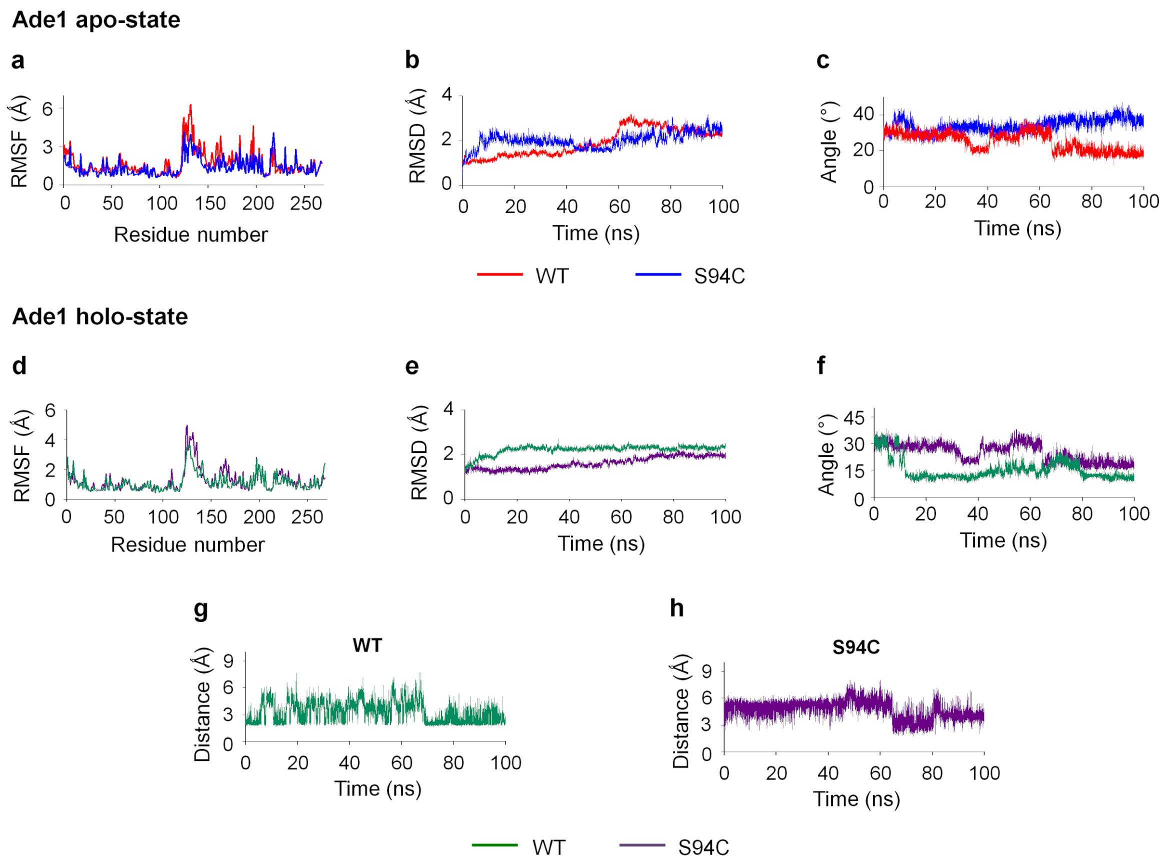

**Figure S16. Structural analysis of Ade1$_{S94C}$ in its apo- and holo-state.** Molecular dynamics simulation explains the loss of enzymatic activity of Ade1$_{S94C}$. **(a)** Side chain root-mean-square fluctuations (RMSF) of Ade1 and Ade1$_{S94C}$ apo-states (red and blue colors, respectively). **(b)** Backbone root-mean-square fluctuations (RMSD) of Ade1 and Ade1$_{S94C}$ apo-states (red and blue colors, respectively). **(c)** Closing angle (θ) was calculated using the coordinates formed by the main chain oxygen atoms of the V127, L27, and M194 residues. Closing angles of Ade1 and Ade1$_{S94C}$ holo-states are colored in red and blue colors, respectively. **(d)** Side chain root-mean-square fluctuations (RMSF) of Ade1 and Ade1$_{S94C}$ holo-states (green and purple colors, respectively). **(e)** Backbone root-mean-square fluctuations (RMSD) of Ade1 and Ade1$_{S94C}$ holo-states (red and blue colors, respectively). **(f)** Closing angle of cap domain. The change from S94 to C94 prejudices the closing of the cap domain. Average distance between the pairs **(g)** S94/H245 and **(h)** C94/H245 increased from ~3 to ~5.5 Å, respectively.

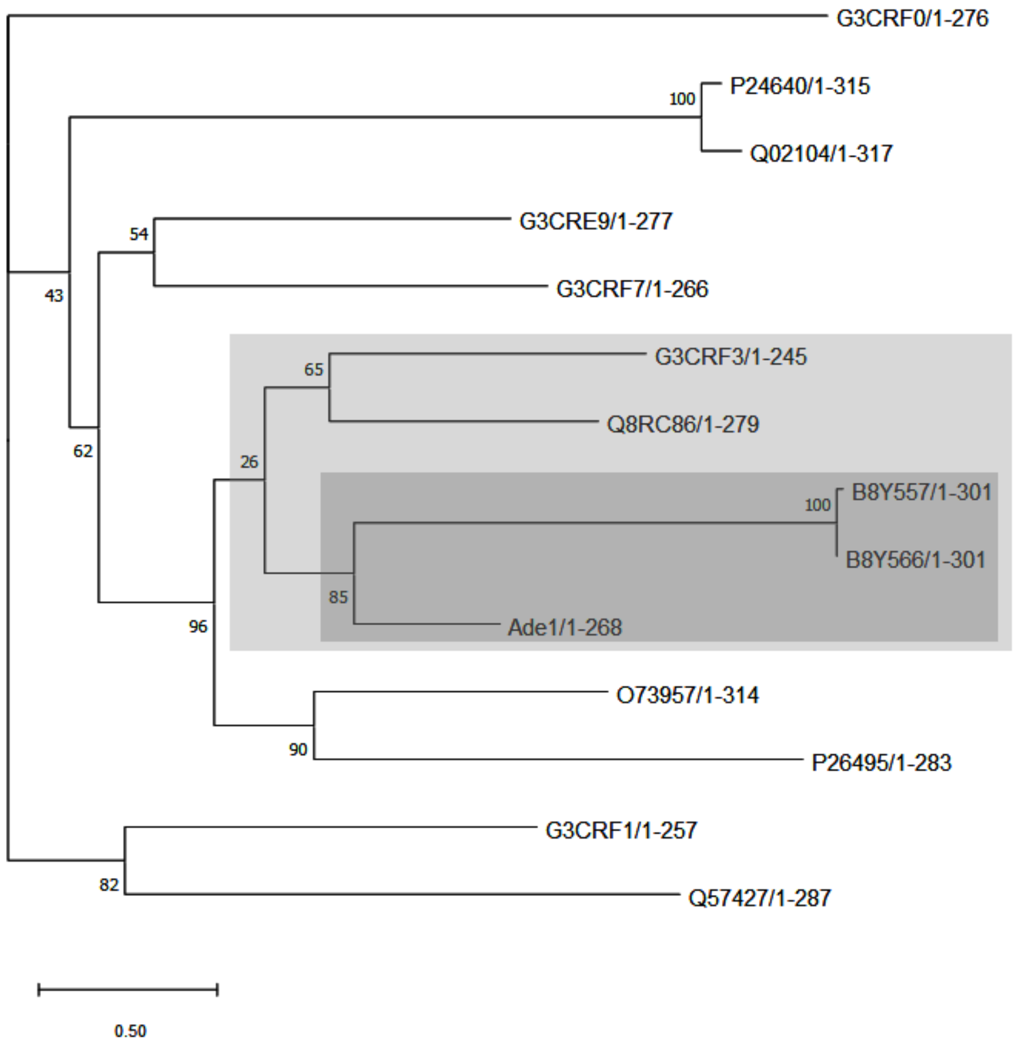


**Figure S17. Dendrogram of the family V of lipolytic enzymes.** Sequences of members belonging to family V were used to build the dendrogram representation. It is used to observe the similarity between lipolytic enzymes used in **Figure 1b**. Gray color box indiques the protein sequences with higher sequence similarity to Ade1. Phylogenetic tree was built using a neighbor-joining algorithm. Gray color more intense shows sequences more similar to Ade1.

### Tables

**Table S1.** Nucleic acid sequence as well as Ade1 amino acid sequence.

| *ade1* nucleic acid sequence | *bp* |
|---|---|
| ATGCTATATGCTCAGGTCAACGGCATCAATTTGCACTATGAGATCGAAGGTCAGG<br>GGCAGCCTCTGTTACTCATTATGGGGTTAGGTGCGCCAGCGGCAGCCTGGGAC<br>CCAATATTCGTCCAGACGCTGACCAAGACTCATCAAGTCATCATTTACGACAACC<br>GCGGCACAGGACTCTCGGACAAGCCTGATATGCCCTATTCGATCGCTATGTTCG<br>CCAGCGATGCTGTTGGCTTGCTTGATGCACTCAATATCCCTCGAGCCCACGTCTT<br>TGGCGTATCAATGGGTGGGATGATCGCTCAGGAGCTGGCGATTCACTACCCGCA<br>GCGAGTCGCCAGCTTGATTCTGGGTTGCACGACACCTGGCGGTAAGCATGCGGT<br>CCCGGCTCCACCAGAGTCGCTGAAAGCTTTAGAAGGTCGAGCTGGACTAACTCC<br>GGAAGAGGCAATTCGGGAAGGTTGGAAACTCTCTTTTTCTGAGGAATTTATCCAC<br>ACGCACAAGGCTGAACTGGAAGCACACATACCGCGGCTGCTTGCCCAACTTACC<br>CCACGCTTTGCCTATGAGCGCCACTTCCAGGCAACTATGACACTGAGAGTTTTCA<br>AGCAGCTCAAAGAGATTCAAGCGCCAACCTTGGTAGCGACAGGACGCGATGACA<br>TGCTCATCCCCGCAGTAAATTCCGAGATTCTGGCGCGCGAGATTCCCGGTGCCG<br>AGTTGGCCATCTTCGAAAGCGCCGGCCACGGGTTCGTGACCTCGGCGCGTGAG<br>CCGTTCCTCAAAGTGTTGAAGGAGTTTCTGGCACGGCAAAGCGTGTAGGCATAG<br>CAAGCCAGCGTCCAGAGTCTAGA | 804 |

| Ade1 amino acid sequence | aa |
|---|---|
| MLYAQVNGINLHYEIEGQGQPLLLIMGLGAPAAAWDPIFVQTLTKTHQVIIYDNRGTGL<br>SDKPDMPYSIAMFASDAVGLLDALNIPRAHVFGVSMGGMIAQELAIHYPQRVASLILG<br>CTTPGGKHAVPAPPESLKALEGRAGLTPEEAIREGWKLSFSEEFIHTHKAELEAHIPR<br>LLAQLTPRFAYERHFQATMTLRVFKQLKEIQAPTLVATGRDDMLIPAVNSEILAREIPG<br>AELAIFESAGHGFVTSAREPFLKVLKEFLARQSV | 268 |

**Table S2. Oligonucleotides used in the cloning.** Underlines mark restriction sites and red nucleotides mark mutation sites. The symbols Fw and Rv are Forward and Reverse, respectively.

| | |
|---|---|
| **Cloning primers** | |
| Ade1 | Sequence 5' - 3' |
| Ade1F | TTTTTCATATGCTATATGCTCAGGTCAACGGC |
| Ade1R | TTTTGGATCCCTACACGCTTTGCCGGT |
| **Site-specific mutagenesis primers** | |
| S94A - Fw | CCACGTCTTTGGCGTAGCAATGGGTGGGATGATCGCTCA |
| S94A - Rv | TGAGCGATCATCCCACCCATTGCTACGCCAAAGACGTGG |
| S94C - Fw | CCACGTCTTTGGCGTATGCATGGGTGGGATGATCGCTCA |
| S94C - Rv | TGAGCGATCATCCCACCCATGCATACGCCAAAGACGTGG |

**Table S3.** X-ray data collection and refinement statistics for Ade1 in the presence of 4mM tributyrin. There are 4 molecules in the asymmetric unit. Ade1 structure was deposited in the protein data bank with PDB ID 6EB3.

| | |
|---|---|
| **X-ray data collection** | |
| Space group | $P2_12_12$ |
| Unit-cell parameters (Å) | |
| a (Å) | 71.7 |
| b (Å) | 106.0 |
| c (Å) | 146.0 |
| Resolution range (Å)[1] | 30.00 – 2.30 (2.38 - 2.30) |
| $N_{0.}$ observed reflections<br>$N_{0.}$ unique reflections | 296,448<br>50,655 |
| <I/errorI>[1] | 13.4 (2.3) |
| Multiplicity[1] | 5.7 (5.9) |
| Completeness (%)[1] | 99.9 (100.0) |
| Rmerge (%)[1,2] | 4.3 (36.9) |
| λ (Å) | 1.54 |
| **Model Refinement** | |
| Resolution range (Å) | 30.00 – 2.30 |
| $R_{factor}/R_{free}$ (%) | 19.3/23.3 |
| No. reflections | 57,040 |
| No. of water molecules | 391 |
| **Stereochemistry** | |
| r.m.s deviation from ideal geometry | |
| Bond bond (Å) | 0.012 |
| Bond angle (°) | 1.549 |
| Protein average B-factors (Å$^2$) | 43.4 |
| **Ramachandran plot** (%)[3] | |
| Most favored regions | 96.9 |
| Allowed regions | 3.1 |
| Disallowed regions | 0.0 |

[1] Values in parentheses refer to the highest resolution shell .

[2] $R_{merge} = (\Sigma | I-\langle I\rangle |) / \Sigma(I)$

$R_{factor} = \Sigma | F_{obs} - F_{calc} | / \Sigma | F_{obs} |$

$R_{free}$ was calculated using 5.1 % of the reflections selected randomly and omitted from the refinement.

[3]Calculated in PROCHECK from the ccp4i interface

[4]Calculated from the PISA web server (http://www.ebi.ac.uk/pdbe/pisa/)

**Table S4.** Analysis of three-dimensional structural similarities between the Ade1 structure and others deposited in public databases using DALI web-based service. The search for structural similarities used as input was the Ade1 chain B. The Table describes the main results. These results show that Ade1 shares three-dimensional structural similarities with a wide range of α/β-hydrolases members that have different enzymatic functions.

| PDB ID-Chain | Z-score | RMSD (Å) | id % | Enzyme name or short description |
|---|---|---|---|---|
| 2xua-H | 34.1 | 2.3 | 27 | 3-OXOADIPATE ENOL-LACTONASE |
| 3fob-A | 31.6 | 2.4 | 22 | BROMOPEROXIDASE |
| 1u2e-A | 31.2 | 2.0 | 26 | 2-HYDROXY-6-KETONONA-2,4-DIENEDIOIC ACID |
| 1va4-A | 31.0 | 2.4 | 24 | ARYLESTERASE |
| 4uhc-A | 31.0 | 2.8 | 25 | ESTERASE |
| 1q0r-A | 30.9 | 2.3 | 27 | ACLACINOMYCIN METHYLESTERASE |
| 4dgq-A | 30.8 | 2.4 | 20 | NON-HEME CHLOROPEROXIDASE |
| 4lxh-A | 30.4 | 2.3 | 23 | MCP HYDROLASE |
| 1c4x-A | 30.2 | 2.4 | 21 | (2-HYDROXY-6-OXO-6-PHENYLHEXA-2,4-DIENOAT |
| 4opm-A | 29.6 | 2.3 | 23 | LIPASE |
| 5ng7-A | 28.9 | 2.1 | 24 | EPOXIDE HYDROLASE |
| 2vf2-A | 28.7 | 2.5 | 24 | 2-HYDROXY-6-OXO-6-PHENYLHEXA-2,4-DIENOATE |
| 3vvl-B | 28.3 | 2.9 | 23 | HOMOSERINE O-ACETYLTRANSFERASE |
| 3kxp-A | 28.0 | 2.3 | 23 | ALPHA-(N-ACETYLAMINOMETHYLENE)SUCCINIC ACID |
| 5frd-A | 27.8 | 2.9 | 23 | CARBOXYLESTERASE (EST-2) |
| 1wom-A | 27.5 | 3.0 | 18 | SIGMA FACTOR SIGB REGULATION PROTEIN RSBQ |
| 2oci-A | 27.4 | 2.3 | 23 | VALACYCLOVIR HYDROLASE |
| 3nwo-A | 27.3 | 2.6 | 22 | PROLINE IMINOPEPTIDASE |
| 1y37-A | 27.2 | 2.6 | 21 | FLUOROACETATE DEHALOGENASE |
| 3kda-A | 26.4 | 2.7 | 18 | CFTR INHIBITORY FACTOR (CIF) |
| 4l0c-A | 26.3 | 2.4 | 25 | DEFORMYLASE |

| | | | | |
|---|---|---|---|---|
| 6f9o-A | 25.9 | 2.8 | 22 | HALOALKANE DEHALOGENASE |
| 6eic-B | 25.4 | 2.6 | 19 | MYCOBACTERIUM TUBERCULOSIS MONOGLYCERIDE LIPASE |
| 6a9d-A | 25.3 | 3.0 | 17 | HYPOSENSITIVE TO LIGHT 7 |
| 2vav-B | 25.2 | 3.1 | 20 | ACETYL-COA--DEACETYLCEPHALOSPORIN C ACETYLTRANSFE |
| 5xmw-A | 24.6 | 2.8 | 18 | ZEARALENONE LACTONASE |
| 6ny9-B | 23.6 | 2.6 | 18 | MYCOPHENOLIC ACID ACYL-GLUCURONIDE ESTERASE |
| 2psf-A | 23.3 | 2.9 | 15 | RENILLA-LUCIFERIN 2-MONOOXYGENASE |
| 4gdm-C | 23.2 | 3.3 | 15 | 2-SUCCINYL-6-HYDROXY-2,4-CYCLOHEXADIENE-1-CARBOXY |
| 3qit-B | 23.1 | 2.4 | 19 | POLYKETIDE SYNTHASE |
| 4qlo-A | 23.1 | 3.6 | 16 | HOMOSERINE O-ACETYLTRANSFERASE |
| 3pf8-A | 22.6 | 2.7 | 17 | CINNAMOYL ESTERASE |
| 6ix4-B | 22.2 | 2.9 | 13 | MICROSOMAL EPOXIDE HYDROLASE |
| 2wm2-C | 21.9 | 3.2 | 16 | 1-H-3-HYDROXY-4-OXOQUINALDINE 2,4-DIOXYGENASE |
| 1imj-A | 21.5 | 2.1 | 19 | CCG1-INTERACTING FACTOR B |
| 2o2g-A | 21.3 | 2.3 | 21 | DIENELACTONE HYDROLASE |
| 3fnb-A | 20.9 | 2.8 | 14 | ACYLAMINOACYL PEPTIDASE SMU_737 |
| 2jbw-B | 20.7 | 2.6 | 16 | 2,6-DIHYDROXY-PSEUDO-OXYNICOTINE HYDROLASE |
| 6vap-A | 20.6 | 3.2 | 18 | THIOESTERASE |
| 6ba8-A | 20.4 | 3.2 | 14 | IRON AQUISITION YERSINIABACTIN SYNTHESIS ENZYME, |
| 3o4j-C | 20.3 | 2.8 | 15 | ACYLAMINO-ACID-RELEASING ENZYME |
| 1ivy-A | 20.0 | 3.0 | 15 | HUMAN PROTECTIVE PROTEIN |
| 4ao8-A | 19.6 | 2.4 | 16 | ESTERASE |
| 5y57-C | 19.4 | 3.0 | 16 | PYRETHROID HYDROLASE |
| 3wwp-A | 19.1 | 3.1 | 15 | (S)-HYDROXYNITRILE LYASE |
| 5cml-A | 19.0 | 2.9 | 18 | OSMC FAMILY PROTEIN |

| | | | | |
|---|---|---|---|---|
| 1auo-A | 18.7 | 2.5 | 16 | CARBOXYLESTERASE |
| 3n2z-B | 18.7 | 3.2 | 13 | LYSOSOMAL PRO-X CARBOXYPEPTIDASE |
| 3hxk-A | 18.6 | 2.7 | 14 | SUGAR HYDROLASE |
| 4fhz-A | 18.2 | 2.7 | 19 | PHOSPHOLIPASE/CARBOXYLE STERASE |
| 4kry-E | 18.1 | 2.5 | 17 | ACETYL ESTERASE |
| 2cb9-A | 18.0 | 3.0 | 11 | FENGYCIN SYNTHETASE |
| 6kd0-A | 17.5 | 2.8 | 12 | VIBRALACTONE CYCLASE |
| 1lzk-A | 17.4 | 2.9 | 18 | HEROIN ESTERASE |
| 3hlk-B | 17.3 | 2.4 | 11 | ACYL-COENZYME A THIOESTERASE 2, MITOCHONDRIAL |
| 2h7y-A | 16.6 | 3.0 | 12 | TYPE I POLYKETIDE SYNTHASE PIKAIV |
| 1jkm-B | 16.4 | 2.8 | 14 | BREFELDIN A ESTERASE |
| 1jjf-A | 16.1 | 2.6 | 13 | ENDO-1,4-BETA-XYLANASE Z |
| 3i2g-A | 15.9 | 2.8 | 17 | COCAINE ESTERASE |
| 3iii-A | 15.9 | 2.9 | 12 | COCE/NOND FAMILY HYDROLASE |
| 4xjv-A | 15.9 | 2.8 | 15 | S-ACYL FATTY ACID SYNTHASE THIOESTERASE, MEDIUM C |
| 3c6b-A | 15.7 | 2.8 | 17 | S-FORMYLGLUTATHIONE HYDROLASE |
| 3f98-A | 15.5 | 2.7 | 14 | PLATELET-ACTIVATING FACTOR ACETYLHYDROLASE |
| 3wl5-A | 15.5 | 3.0 | 10 | OXIDIZED POLYVINYL ALCOHOL HYDROLASE |
| 4j0d-A | 15.5 | 3.3 | 14 | TANNASE |
| 3ils-A | 15.4 | 3.3 | 13 | AFLATOXIN BIOSYNTHESIS POLYKETIDE SYNTHASE |
| 4e15-B | 15.2 | 3.0 | 13 | KYNURENINE FORMAMIDASE |
| 6rt8-A | 15.1 | 2.9 | 12 | CATHARANTHINE SYNTHASE |
| 3og9-A | 15.0 | 3.2 | 15 | PROTEIN YAHD A COPPER INDUCIBLE HYDROLASE |
| 6rs4-B | 14.8 | 3.0 | 11 | TABERSONINE SYNTHASE |
| 1cle-A | 14.4 | 2.9 | 10 | CHOLESTEROL ESTERASE |
| 6eop-A | 14.4 | 2.8 | 19 | DIPEPTIDYL PEPTIDASE 8 |
| 3vkf-A | 14.1 | 2.8 | 14 | NEUROLIGIN-1 |
| 1mx9-D | 13.7 | 3.6 | 18 | LIVER CARBOXYLESTERASE I |
| 5ie4-C | 13.5 | 2.6 | 22 | ZEARALENONE HYDROLASE |

| | | | | |
|---|---|---|---|---|
| 4be4-A | 13.0 | 3.0 | 11 | STEROL ESTERASE |
| 5nuu-A | 12.7 | 2.8 | 13 | ACETYLCHOLINESTERASE |
| 4qnn-A | 11.1 | 3.6 | 13 | PHOSPHOLIPASE |
| 2czq-A | 10.8 | 2.9 | 13 | CUTINASE-LIKE PROTEIN |
| 4uyz-D | 10.0 | 3.5 | 9 | PROTEIN NOTUM HOMOLOG |
| 1bs9-A | 9.5 | 3.4 | 13 | ACETYL XYLAN ESTERASE |
| 2xa2-A | 6.9 | 3.4 | 11 | TREHALOSE-SYNTHASE TRET |

**Table S5. Quorum-quenching activity of different enzymes.** The table has information about the enzyme, organism, biological function and if there is the presence or absence of metal during the enzymatic catalysis. There are proteins that make quorum-quenching activity.

| Name | Organism | Function | metal/no metal |
|---|---|---|---|
| 3-hydroxy-palmitic acid methyl ester hydrolases | metagenomic library | esterase[32] | no metal reported |
| quorum sensing molecule 3OH-PAME | XB7 and XB122 (*Pseudomonas aeruginosa*) and XB102 (*Stenotrophomonas maltophilia*) | esterase[33] | $Cu^{2+}$ and $Ca^{2+}$ enhanced activity |
| Est816 esterase | Turban Basin metagenomic library | esterase[34] | no metal reported |
| Porcine kidney acylase (PKA), Porcine liver esterase(PLE) and horse liver esterase (HLE) | Porcine and Horse | acylase and esterase[35] | no metal reported |
| Est816 esterase | Turban Basin metagenomic library | esterase[36] | slightly promoted by $Ca^{2+}$ |
| QsdH | *Pseudoalteromonas byunsanensis* | esterase[37] | at 0.2 mM slight enhances activity ($Zn^{2+}$, $Ni^{2+}$, $Cu^{2+}$, $Ba^{2+}$, $Mg^{2+}$, $Sr^{2+}$, $Ca^{2+}$ and $Mg^{2+}$), at 10 mM decreases |

| | | | |
|---|---|---|---|
| | | | the activity |
| beta-hydroxypalmitate methyl ester hydrolase | *Ideonella* sp. 0-0013 | esterase[38] | $Fe^{2+}$ and $Sr^{2+}$ inhibit the enzyme activity. $Na^{+}$ and $K^{+}$ promoted enzymatic activity. $Zn^{2+}$ and $Mg^{2+}$ does not inhibit the enzymatic activity. |
| 3-hydroxy-2-methyl-4(1H)-quinolone 2,4-dioxygenase (Hod) | *Arthrobacter nitroguajacolicus* | dioxygenase[39] | no metal reported |
| AiiA | *Bacillus* sp. strain DMS133 | esterase[40] | no metal reported |
| AiiAS1−5 and EstS1−5 | *Altererythrobacter* sp. S1-5 | esterase[41] | no metal reported |
| AHL-lactonase | *Rhodococcus* sp. BH4 | esterase[42] | no metal reported |
| BpiB05 | metagenome-derived hydrolase | esterase[43] | $Ca^{2+}$ are required for carrying out the catalysis |
| AqdC | *M. abscessus* subsp. abscessus | esterase[44] | no metal reported |
| AiiAQSI-1 | *Bacillus* sp. strain QSI-1 | esterase[45] | no metal reported |
| EstDL30 | Alluvial soil metagenomic library: phylogenetic analysis suggests that is produced by *Bacillus subtilis* (P37967), | esterase[46] | no metal reported |

| | | | |
|---|---|---|---|
| | *Streptomyces coelicolor* A3(2) (CAA22794), and *Arthrobacter oxydans* (Q01470) | | |
| AHL-lactonase | Human, mouse and fish | esterase[47] | Enzyme dependent of $Ca^{2+}$ |
| metallo-beta - lactamase (AiiA) | *Bacillus thuringiensis* | esterase[48] | Enzyme dependent of $Zn^{2+}$. AiiA contains a bimetallic active site coordinated by two Asp residues and five His. |
| VmoLac | *Vulcanisaeta moutnovskia* | esterase[49] | VmoLac is a bi-cobalt metalloenzyme |
| N-acyl-homoserine lactone acylase | *Ralstonia solanacearum* GMI1000 | acylase[50] | no metal reported |
| AidF | *Ochrobactrum intermedium* D-2 | esterase[51] | no metal reported |
| FadY | *Acinetobacter lactucae* strain QL-1 | acyl-CoA synthetase[52] | no metal reported |
| no enzymatic characterization | no microrganism was characterized | acylase[53] | no metal reported |
| GcL | *Geobacillus caldoxylosilyticus* | esterase[54] | no metal reported |
| PFE esterase | *Burkholderia anthina* HN-8 | esterase[55] | no metal reported |

| | | | |
|---|---|---|---|
| PLL(PTE-like lactonase) | *Mycobacterium avium* subsp. paratuberculosis K-10 | esterase[56] | no metal reported |
| QsdA | *Rhodococcus erythropolis* | phosphotriesterases[57] | no metal reported |

**Table S7. Different enzymes with hysteresis behavior.** The table has information about enzymes, transient lag or burst, organism, biological function and if there is the presence or absence of metal during the enzymatic catalysis and/or optimal pH.

| **Enzyme** | **Organism** | **Description** | **Type of transient** | **Metal (optimal pH)** |
|---|---|---|---|---|
| Pyruvate kinase[58] | *Bacillus licheniformis* | ATP obtention (metabolism) | Lag | Activation by $Mg^{2+}$ (7.0 to 7.4) |
| Glutamine phosphoribosyl pyrophosphate amidotransferase[59] | Pigeon liver | Purine biosynthesis (metabolism) | Lag | Activation by $Mg^{2+}$ (8.0) |
| D-Lactate dehydrogenase[60] | *Escherichia coli* | Fermentation – anaerobic metabolic pathway (metabolism) | Lag | (6.4 – 7.5) |
| Hexokinase B[61] | *Saccharomyces cerevisiae* | Regulation of carbon catabolism (Metabolism) | Burst | (7.0 - 8.0) |
| StEH1[62] | *Solanum tuberosum* | Epoxide hydrolase | Burst | Activation by temperatura change (8.0 to 9.0) |
| BChE[63] | Human | Butyrylcholinesterase (metabolism) | Burst | Activation by hydrostatic pressure, temperature, salts and pH |
| Nitrate reductase[64] | *Cucurbita maxima* | Nitrate assimilation (metabolism) | Lag | Activation by phosphorylation state (7.5) |
| Alkaline phosphatase (AP)[65] | Calf intestine | metabolic process | ? | (10.0 to 10.8) |

| | | | | |
|---|---|---|---|---|
| Trehalase[66] | *Artemia salina* | Dormancy embryos | ? | Intracellular pH dependent |
| ProT (Prothrombin)[67] | Mammals | Blood coagulation | Lag | Activation by VWbp (Von Willebrand factor-binding protein) (7.0) |
| PFK (Phosphofructokinase)[68] | Rat myocardium | Glycolysis (metabolism) | Lag | Alkalinization of the myocardium muscle |
| ATP synthase[69] | *Polytomella sp.* | Energy transduction in mitochondria (Oxidative phosphorilation metabolism) | Lag | Temperature (8.0) |
| Oxidized fructose 1,6-bisphosphatase[70] | Chloroplast | Glycolisis | Lag | fructose 2,6-bisphosphate (substrate analog), magnesium (7.5) |
| α-Acetylgalactosaminidase[71] | Bovine | Glycoside hydrolase | Lag | (4.7 - 5.0) |
| Homoserine dehydrogenase[72] | *Escherichia coli* | Synthesis of L-homoserine | Burst | $K^+$ (6.9) |

**Table S8. Hydrogen bonding occupancy (%) between the side chains of catalytic residues (only S94/C94 and H245) and nearby residues in the holo-state.** Hydrogen bond occupancy (%) analysis of the catalytic residues (S94/C94 and H245) that make contact with nearby residues inside the enzymatic cavity along molecular dynamics simulations. Cut-off: hydrogen bonding distance of 3 Å between hydrogen and nitrogen or oxygen.

| HB donor | HB acceptor | Ade1 (%) | Ade1$_{S94C}$ (%) |
|---|---|---|---|
| H245 (Hδ1) | D217 (Oγ1) | 69.3 | 73.8 |
| S94/C94 (Hγ1) | H245 (Nε2) | 13.5 | - |
